# Cortical cell stiffness is independent of substrate mechanics

**DOI:** 10.1101/829614

**Authors:** Johannes Rheinlaender, Andrea Dimitracopoulos, Bernhard Wallmeyer, Nils M. Kronenberg, Kevin J. Chalut, Malte C. Gather, Timo Betz, Guillaume Charras, Kristian Franze

## Abstract

Cortical stiffness is an important cellular property that changes during migration, adhesion, and growth. Previous atomic force microscopy (AFM) indentation measurements of cells cultured on deformable substrates suggested that cells adapt their stiffness to that of their surroundings. Here we show that the force applied by AFM onto cells results in a significant deformation of the underlying substrate if it is softer than the cells. This ‘soft substrate effect’ leads to an underestimation of a cell’s elastic modulus when analyzing data using a standard Hertz model, as confirmed by finite element modelling (FEM) and AFM measurements of calibrated polyacrylamide beads, microglial cells, and fibroblasts. To account for this substrate deformation, we developed the ‘composite cell-substrate model’ (CoCS model). Correcting for the substrate indentation revealed that cortical cell stiffness is largely independent of substrate mechanics, which has significant implications for our interpretation of many physiological and pathological processes.

## Introduction

*In vivo*, cells respond to the mechanical properties of their environment^1,2^. As the stiffness of any tissue critically depends on the mechanical properties of its constituent cells, cell mechanics measurements are key to understanding many complex biological processes. Over the last decades, atomic force microscopy (AFM) has emerged as a gold standard to assess the mechanical properties of cells^3–6^. In AFM measurements, a force is applied to the cell surface, and the resulting deformation is used to calculate an apparent elastic modulus, which is a measure of the cell’s stiffness. Depending on the force applied, different cellular structures contribute differently to the measured elastic moduli^6^. AFM indentation measurements of cells using low stresses (force per area) and thus resulting in small strains (relative deformations) mainly probe peripheral cellular structures including the actomyosin cortex^7^ and the pericellular coat^8^. The measured apparent elastic moduli can then be interpreted as an effective cortical cell stiffness.

Previous AFM studies suggested that the cortical stiffness of cells increases with increasing substrate stiffness^9–13^. The application of blebbistatin, which blocks myosin II function and thus cell contractility, abolished the apparent stiffening of the cells on stiffer substrates. Hence, it was hypothesized that, as cells increase their traction forces on stiffer substrates, the increased pre-stress of the actomyosin network leads to its non-linear stress stiffening and accordingly to an overall stiffening of the cells^9^.

In AFM indentation measurements, the relation between the loading force *F* and the overall sample indentation *δ* is mostly modeled using the Hertz model^14^, which in the case of a spherical probe is as follows:

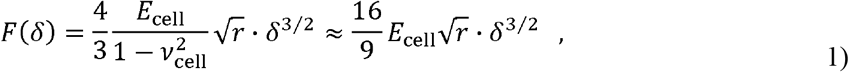

where *r* is the probe radius, *v*_cell_ is the cell’s Poisson’s ratio, which usually is close to *v*_cell_∼0.5^15^, and *E*_cell_ is the apparent elastic modulus of the cell. The only quantities recorded during an experiment are the cantilever’s vertical displacement Δ*z* and its deflection *d*. *d* is used to calculate the applied force *F* = *k* · *d*, where *k* is the cantilever’s spring constant. The indentation depth *δ* = Δ*z* – *d* is inferred from these quantities based on the key assumption that the sample is deformed but not the underlying substrate (Fig. 1a). However, while this condition is clearly fulfilled for cells cultured on glass or tissue culture plastics, it may no longer hold for cells cultured on soft matrices mimicking the mechanical properties of the physiological cell environment^16^.

**Figure 1:**
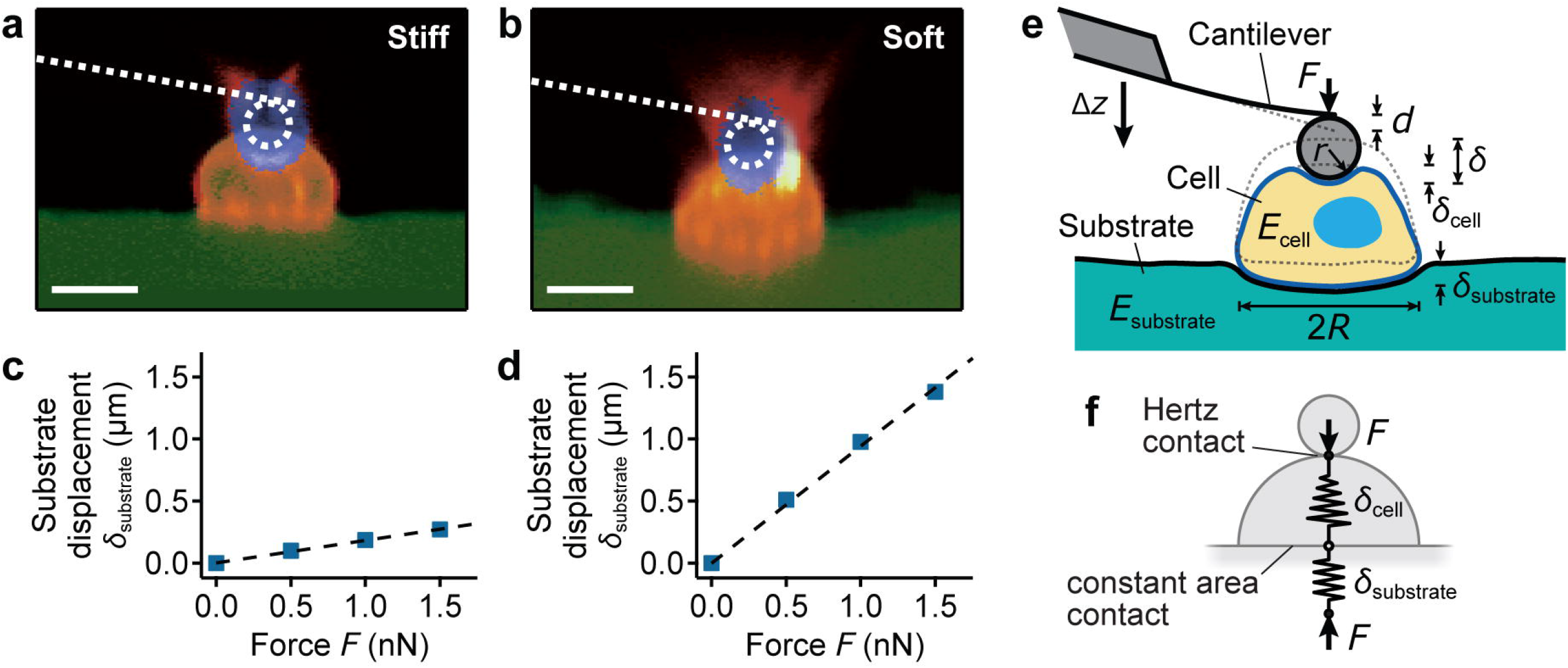
Quantification of substrate displacements in AFM indentation measurements of cells. **(a, b)** Confocal *z-x* profiles of microglial cells (orange) cultured on **(a)** stiff (≈ 2 kPa) and **(b)** soft (≈ 100 Pa) substrates (green). The AFM probe (blue) is applying a loading force of *F* = 1 nN on each cell. Scale bars: 10 μm. **(c, d)** Relationship between substrate displacements obtained from confocal images of the cells shown in **(a)** and **(b)** and the applied force *F* on **(c)** stiff and **(d)** soft substrates (see also Supplementary Fig. 1g-i). Forces exerted on cells by AFM indentation result in significant deformations particularly of soft substrates. **(e)** Schematic of an AFM cantilever with a spherical probe of radius *r* pushing on a cell with elastic modulus *E*_cell_ and radius *R* bound to a substrate with elastic modulus *E*_substrate_. The measured indentation *δ* is a combination of the indentation of the cell, *δ*_cell_, and that of the substrate, *δ*_substrate_. The dotted outline indicates the undeformed state. Δ*z* denotes vertical cantilever displacement, *d* cantilever deflection. **(f)** Schematic of the mechanical system, consisting of the two springs in series, which both experience the same force. The tip-cell contact follows the nonlinear Hertz model^14^, and the cell-substrate contact follows a linear force-indentation relation due to the largely constant contact area, similar to other analytical contact models^19,20^.

## Results

### AFM indentation pushes cells into soft substrates

Indeed, when we cultured microglial cells on polyacrylamide substrates with stiffnesses ranging from *E*_substrate_ = 50 Pa to 20 kPa and probed them by combined AFM/confocal laser scanning microscopy, forces exerted on the cells led to substantial substrate deformations, in contradiction with analytical assumptions (Fig.1, Supplementary Fig. 1). On stiffer substrates (*E*_substrate_ ≈ 2 kPa), force applied by the cantilever on cells resulted in negligible vertical substrate displacements (Fig. 1a). On softer substrates (*E*_substrate_ ≈ 100 Pa), however, applied forces resulted in significant vertical substrate displacements on the order of 1 μm (Fig. 1b). Moreover, the substrate displacement depended linearly on the loading force (Fig. 1c, d, 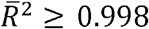), with a small apparent deformability *c* = *δ*_substrate_/*F* of around 0.1 μm/nN for stiff substrates (Fig. 1c) and a significantly larger deformability of *c*∼0.9 μm/nN for soft substrates (Fig. 1d) (see also Supplementary Figs. 1g-i, 2e).

Hence, the indentation *δ* inferred from AFM measurements is actually the sum of the indentation of the cell *δ*_cell_ and that of the underlying substrate *δ*_substrate_, signifying that Δ*z* – *d* = *δ*_cell_ + *δ*_substrate_ (Fig. 1e). On hard substrates, *δ*_substrate_ is negligible and *δ*_cell_ can be directly inferred from the measurements as usually done. However, on soft substrates, excluding *δ*_substrate_ leads to an overestimation of *δ*_cell_ and thus to an underestimation of the cell’s apparent elastic modulus *E*_cell_ when using the standard Hertz model (Equation (1)).

### Analytical model to account for substrate deformation

To address this problem, we first considered a simple analytical model to characterize the deformation of an elastic cell in contact with a deformable substrate, similar as two elastic springs in series (Fig. 1f). The force applied by the cantilever onto the cell is balanced by the elastic deformation of the substrate underneath the cell (*i.e*., the force experienced by the substrate is the same as that exerted onto the cell). To investigate this substrate deformation in more detail, we combined AFM with Elastic Resonator Interference Stress Microscopy (ERISM)^17,18^, which quantifies the vertical deformation of deformable substrates with high spatial resolution (Fig. 2). Both the substrate deformation and the stress were maximum under the cell center, where the cantilever was located, and increased linearly with the applied force (Fig. 2a). Substrate deformation and stress also decayed approximately linearly away from the cell center until reaching zero ∼10 μm away from the cantilever (Fig. 2b). The shape of the substrate deformation and stress distribution did not vary for different applied forces (Fig. 2b).

**Figure 2:**
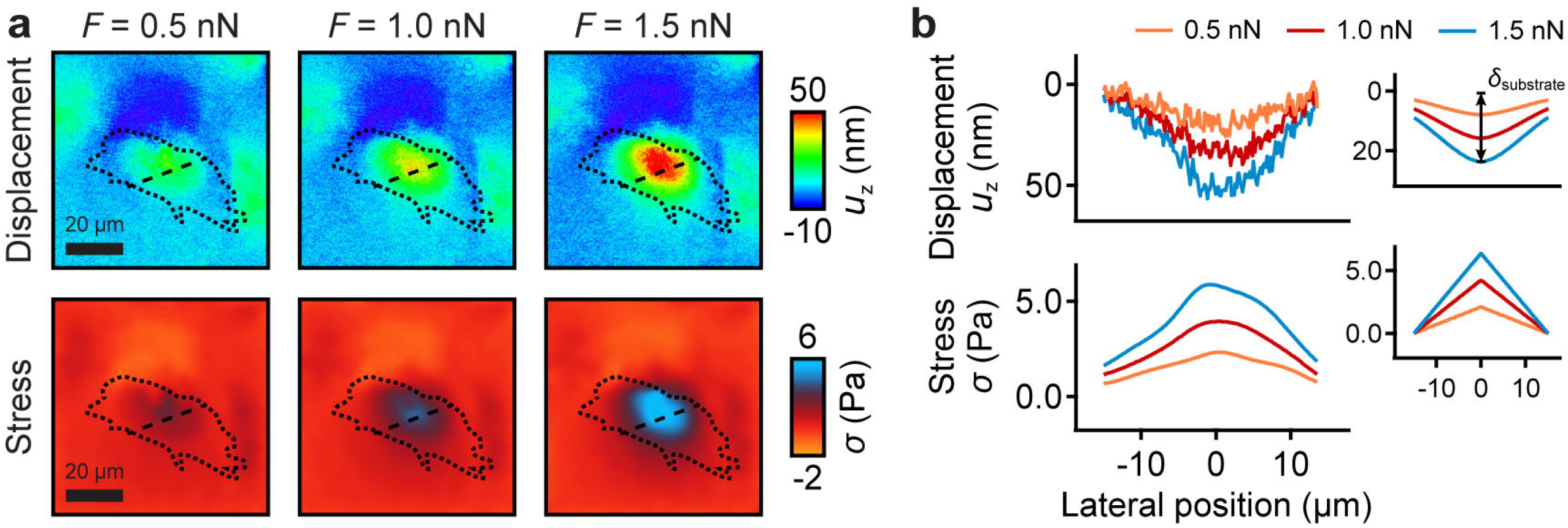
Substrate displacement and stress distribution under cells caused by AFM indentation measurements. **(a)** Displacement (top row) and stress distribution (bottom row) of the substrate measured by ERISM^17^ at different forces *F* applied by AFM. Dotted line: outline of the cell; dashed line: location of profiles shown in (b). **(b)** Profiles of displacement (top) and stress (bottom) under the cell shown in (a). The insets in (b) show displacement (top) and stress (bottom) predicted by the analytical model using an effective cell radius of *R* = 15 μm. There is very good qualitative and quantitative agreement between the model and the data.

We therefore assumed an axisymmetric stress distribution with maximum stress of *σ*_0_ below the cell center and linear decrease from the center to zero within a distance approximated by the cell radius *R* (Fig. 2b). The substrate deformation can then be approximated by the elastic response of a semi-infinite half space due to axisymmetric stress distribution on a circular region^19^, also known as the Boussinesq solution^20^:

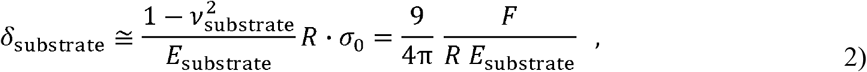

(Fig. 2b) where *v*_substrate_ = 0.5 for polyacrylamide gels^21^. Note that the cell-substrate contact results in a linear force-indentation relation, because the contact area does not change with indentation. As the maximum stress linearly increased with the applied force but its functional form remained unaltered (Fig. 2b), the force-indentation relation will be linear also for any arbitrary cell morphologies.

In contrast, the indentation of the cell follows the non-linear Hertz model^14^:

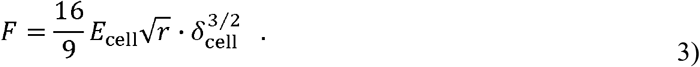

The measured overall indentation *δ* is then a combination of the indentation of the cell and that of the substrate,

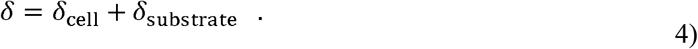

Rearranging Equation (3) and inserting it with Equation (2) into Equation (4) gives the relation between the measured overall indentation and the applied loading force

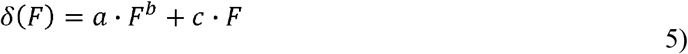

with the pre-factor 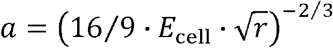, the exponent *b* = 2/3, and

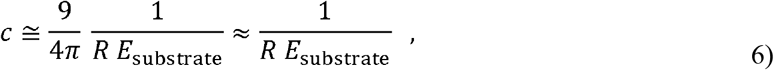

which is proportional to the inverse of the substrate stiffness. *c* can be interpreted as the effective substrate deformability, *i.e*., a measure of how much the substrate (and the cell bottom) is deformed during a measurement.

Since the terms accounting for a cell’s elastic modulus (non-linear Hertz contact) and substrate deformability (linear contact) are linearly independent, fitting Equation (5) to the inverse relationship between force and indentation, with and as free parameters, allows the determination of the cell’s elastic modulus and the substrate deformability independently of each other. We termed this approach the ‘composite cell-substrate model’ (‘CoCS model’). The CoCS model can easily be adapted to other Poisson’s ratios or tip geometries (for example for conical/pyramidal tips using *b* = 1/2 and a different relation for *a* according to the respective contact model^22,23^), and that the tip geometry only affects *δ*_cell_.

### Numerical validation using finite element modeling

To test the effect of substrate stiffness on the measured cell stiffness in AFM experiments, we first used a finite element model (FEM) to generate ground-truth force-distance curves for different ratios of *E*_substrate_/*E*_cell_ and different indenter geometries (Fig. 3, see Methods and Supplementary Figs. 3, 4 for details). We chose a half-spherical geometry to represent the cell (Fig. 1a, b).

**Figure 3:**
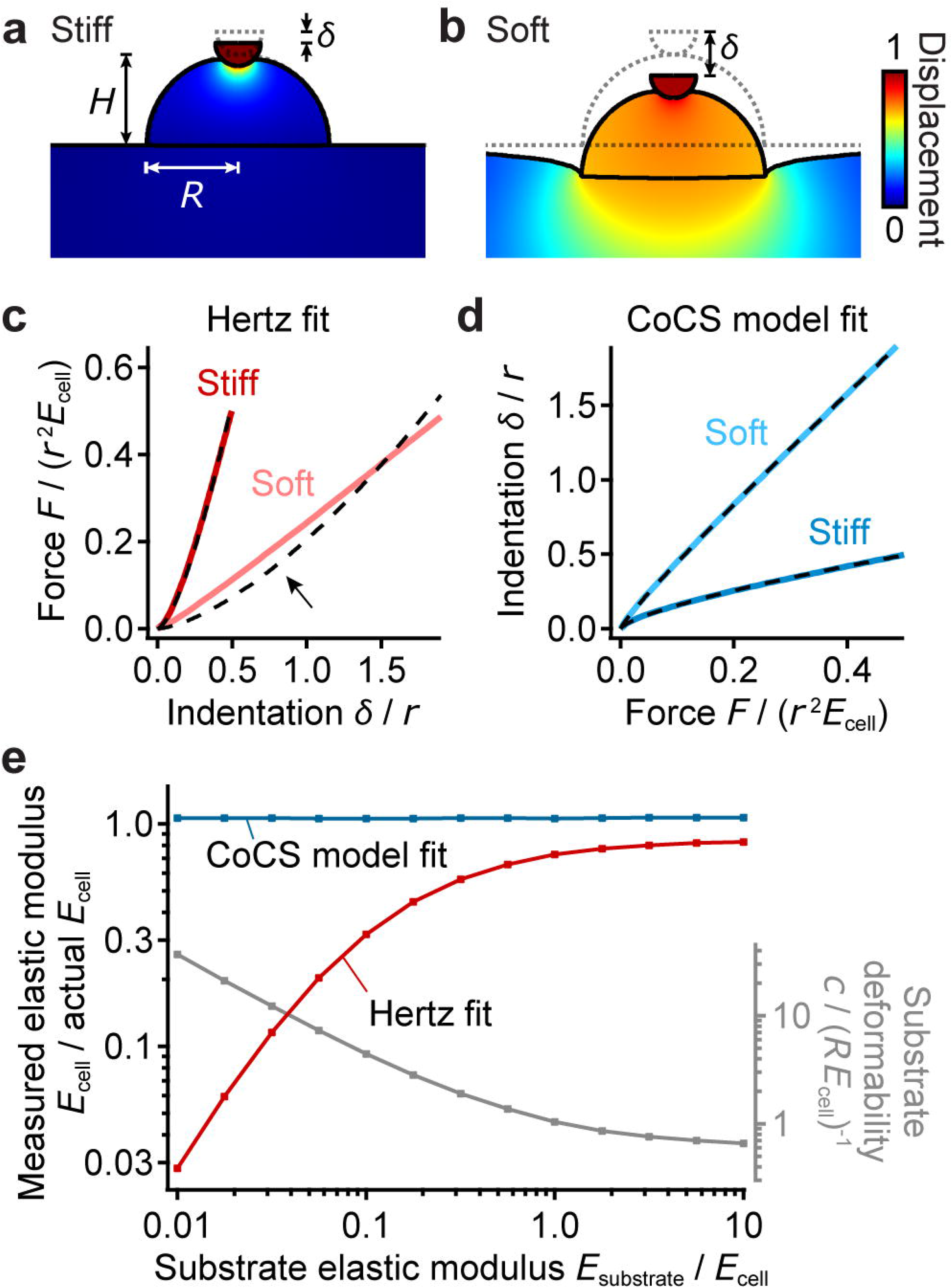
Numerical validation. **(a, b)** Representative FEM results for cells on **(a)** a stiff substrate and **(b)** a soft substrate for the force *F* = 0.5 *r*^2^ *E*_cell_. Color shows material displacement in units of tip displacement. *δ* indicates the measured total indentation relative to the undeformed state (dotted outlines). **(c)** Force *F vs*. indentation *δ* (scaled in units of cell stiffness and tip radius) for cells on stiff and soft substrates analyzed with standard Hertz model fits, Equation (1) (dashed traces). The Hertz model deviates from the data in measurements on soft substrates (arrow). **(d)** Indentation *δ vs*. force *F* for cells on soft and stiff substrates with CoCS model fits, Equation (5) (dashed traces). **(e)** Measured elastic moduli *E*_cell_ in units of the actual elastic moduli of the cells as a function of relative substrate stiffness as obtained fitting force-indentation curves simulated by FEM using a standard Hertz fit (Equation (1), red trace), or using the CoCS model fit (Equation (5), blue trace). Right axis shows substrate deformability obtained from the CoCS model fit. **(a-d)** Parameters of calculations shown: cell height and radius *H* = *R* = 4*r*, *E*_substrate_/*E*_cell_ = 3 (“stiff”) and 0.03 (“soft”).

When we fit the simulated force-distance curves with the Hertz model, the calculated values of *E*_cell_ matched the actual values only on stiff but not on soft substrates, where the cells appeared much softer than they were (0.1 *E*_cell_) and the analytical fit deviated from the simulated curves (Fig. 3c, arrow; 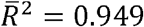 for soft *vs*. 0.999 for stiff substrates). Moreover, while on stiff substrates the force-distance curves always followed the expected *δ*^3/2^-dependency of the Hertz contact, on soft substrates they followed the *δ*^3/2^-dependency only for small forces but approached a linear *δ*-dependency for large forces due to the increasing influence of the substrate deformation (Supplementary Fig. 3b). Hence, the classic Hertz model fit provided the correct cell stiffness only when the substrate stiffness was large compared to the cell stiffness, *E*_substrate_/*E*_cell_ ≫ 1, and it significantly underestimated the cell stiffness when it was comparable to or larger than the substrate stiffness, *E*_substrate_/*E*_cell_ ≲ 1 (Fig. 3e). Similar results were obtained for other tip shapes (Supplementary Fig. 4) and other cell sizes and shapes such as more spherical or well-spread cells (Supplementary Fig. 5).

In contrast, fitting Equation (5) to the same data, plotted as indentation *vs*. force (Fig. 3d), returned correct mechanical properties of the cells irrespective of substrate stiffness (Fig. 3e). The measured cell elastic moduli were now similar on soft and stiff substrates and close to the actual values (here 1.07 *E*_cell_ on stiff and 1.06 *E*_cell_ on soft substrates; Fig. 3d), and the analytical fits matched the simulated curves very well 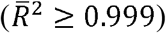. In addition to the real cell elastic moduli, the fits also returned the substrate deformability. The shape of the displacement profile at the cell-substrate interface did not vary for different applied forces (Supplementary Fig. 3c, d), and the substrate deformation linearly depended on the loading force, as well-predicted from the CoCS model fit (Supplementary Fig. 3e).

### Experimental validation using polyacrylamide beads

To test whether the new analysis can accurately determine the stiffness of a real sample with known properties supported by a deformable substrate, we measured elastic polyacrylamide beads of similar diameters as cells with a typical stiffness of *E_bead_* ≅ 1 – 2 kPa by AFM. The mechanical properties of the beads should be independent of the properties of the substrate.

On stiff substrates (*E*_substrate_ ≈ 10 kPa), both the Hertz and the CoCS models fitted the data well (Fig. 4). On soft substrates (*E*_substrate_ ≈ 1 kPa), however, the standard Hertz fit strongly deviated from the experimental force indentation curves (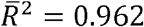 *vs*. 0.999 on stiff substrates), whereas the CoCS model fit still matched the data well 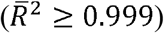 (Fig. 4, Supplementary Fig. 6a). Accordingly, the CoCS model fit estimated bead elastic moduli of around ∼1.2 kPa regardless of the substrate stiffness, while the Hertz model only yielded similar elastic moduli of ∼1.2 kPa on stiff substrates but returned a significantly lower average *E*_bead_ = 0.7 kPa on soft substrates (Fig. 4f).

**Figure 4:**
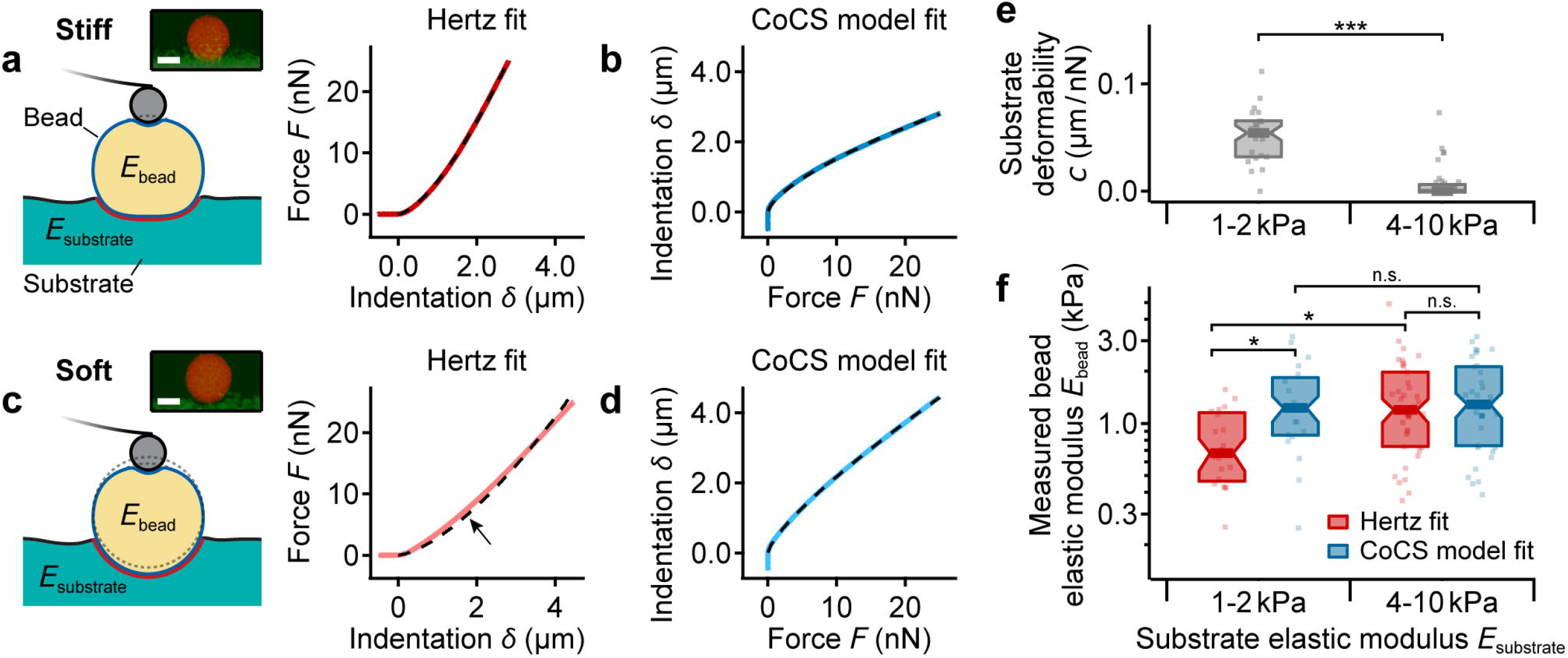
Experimental validation using PAA beads. **(a)** Schematic of AFM measurement of an elastic bead with stiffness *E*_bead_ on a stiff substrate (*E*_substrate_ ≈ 10 kPa ≫ *E*_bead_) and measured force *F vs*. indentation *δ* curve analyzed with standard Hertz model fit, Equation (1). Red solid line: experimental data; dashed trace: fit. **(b)** Same data as in **(a)**, indentation *δ vs*. force *F* analyzed with the CoCS model fit, Equation (5) (dashed trace). **(c)** Schematic for bead on a soft substrate (*E*_substrate_ ≈ 1 kPa ≈ *E*_bead_) and measured force *F vs*. indentation *δ* curve with standard Hertz model fit, Equation (1) (dashed trace). Note the deviation of the model from the experimental data (arrow). **(d)** Same data as in **(c)**, indentation *δ vs*. force *F* with CoCS model fit, Equation (5). Blue solid line: experimental data; dashed trace: fit. The insets in **(a)** and **(c)** show confocal *z-x* profiles of beads (orange) on stiff and soft substrates (green). Scale bar: 10 μm. **(e)** Substrate deformability obtained from CoCS model fits with significantly higher deformability of soft compared to stiff substrates (*P* = 0.012, Wilcoxon-Mann-Whitney *U* test). **(f)** Measured elastic moduli of beads on substrates of different stiffness obtained from Hertz fits (red) and CoCS model fits (blue). Note that the measured bead stiffness is independent of substrate stiffness when using the CoCS model fit (*P* = 0.97, Tukey test), as expected, but significantly depends on the substrate stiffness when using standard Hertz fits (*P* = 0.008, Tukey test). While both models performed similarly well on stiff substrates (*P* = 0.95, Tukey test), on soft substrates the Hertz model yielded significantly lower bead elastic moduli when compared to the CoCS model fit (*P* = 0.03, Tukey test). Box plots: median (band), quartiles (box), standard error (notches), data points (dots); number of beads *N* = 21 and 39 for soft and stiff substrates, respectively. \**P* < 0.05, \*\**P* < 0.01.

The substrate deformability derived from the CoCS model was significantly larger for the soft substrates (*c* = 51 ± 7 nm/nN, average ± SEM) compared to the stiff substrates (*c* = 7.2 ± 1.1 nm/nN) (Fig. 4e), indicating that a significant part of the measured overall indentation on soft but not on stiff substrates was due to the indentation of the substrate by the polyacrylamide beads. Together, these data confirmed that the CoCS model accurately analyzes the elastic stiffness of a sample irrespective of the stiffness of the underlying substrate, in contrast to the commonly used Hertz model, which underestimated sample stiffness on soft substrates.

### Cell stiffness is independent of substrate mechanics

Having validated the ability of our new approach to accurately determine the stiffness of samples regardless of substrate stiffness, we sought to determine if cells indeed adjusted their stiffness to that of their environment^9–13^. As in the bead experiments (Fig. 4), Hertz model fits deviated considerably from the experimental force-distance curves for primary microglial cells cultured on soft matrices (Fig. 5b, 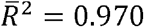) but not on stiff substrates (Fig. 5a, 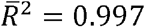). In contrast, the CoCS model fitted both conditions equally well (Fig. 5c, d; 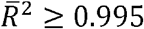) (Supplementary Fig. 6b). The apparent deformability of the substrates significantly increased with decreasing substrate stiffness (Fig. 5e), confirming that a significant part of the overall indentation measured when applying forces to cells cultured on soft substrates originated from the deformation of the substrate.

**Figure 5:**
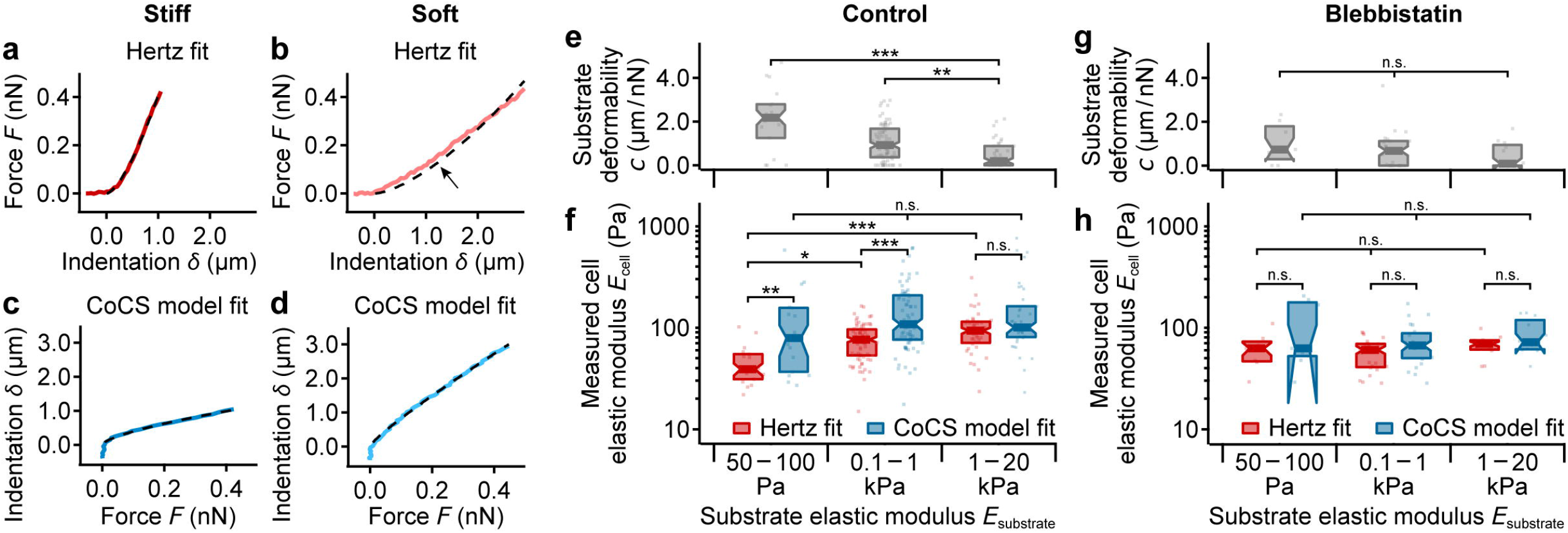
Application to primary microglial cells. **(a, b)** Force *F vs*. indentation *δ* curves for cells on **(a)** stiff and **(b)** soft substrates with Hertz fits (dashed traces). As with beads (Fig. 4), the Hertz model deviated from the experimental data when applied to cells grown on soft substrates (arrow). **(c, d)** Same data as in **(a, b)**, indentation *δ vs*. force *F* on **(c)** stiff and **(d)** soft substrates with CoCS model fit (dashed traces). **(e)** Substrate deformability obtained from CoCS model fits increased significantly with decreasing substrate stiffness (*P* ≤ 0.004, Kruskal Wallis ANOVA followed by Dunn-Holland-Wolfe posthoc test). **(f)** Apparent elastic moduli of live microglial cells on substrates of different stiffnesses as obtained from standard Hertz (red) and CoCS model fits (blue). Similarly to PAA beads, cells were significantly softer on soft and intermediate substrates when analyzed using standard Hertz fits (*P* = 0.03 and *P* = 0.0006, respectively, Tukey posthoc tests), but not when analyzed using the CoCS model fit (*P* = 0.25, one-way ANOVA). Compared to the Hertz model, the CoCS model yielded significantly larger cell stiffnesses on soft and intermediate substrates (*P* = 0.0095 and 4.0 × 10^−6^, respectively, Tukey test), but a similar cell stiffness on stiff substrates (*P* = 0.25, Tukey test). **(g)** Substrate deformability and **(h)** apparent elastic moduli of microglial cells after treatment with the myosin-inhibitor blebbistatin on substrates of different stiffness. Deformability was similar on all substrates (*P* = 0.6, Kruskal-Wallis ANOVA); the measured cell stiffness neither depend on substrate stiffness (*P* ≥ 0.3, one way ANOVA) nor on the fit model (*P* ≥ 0.1, one way ANOVA followed by Tukey posthoc test). Box plots: median (band), quartiles (box), standard error (notches), data points (dots); number of cells (e,f) *N* = 17, 74, and 39 and (g,h) *N* = 7, 24, and 12 for the soft, intermediate, and stiff substrates, respectively. * *P* < 0.05, ** *P* < 0.01, *** *P* < 0.001.

When analyzed using the standard Hertz model, apparent elastic moduli of microglial cells *E*_cell_ decreased significantly from ∼100 Pa on stiffer substrates to ∼40 Pa on soft substrates (Fig. 5f), consistent with previous reports for other cell types^9–13^ (see also Fig. 6). In contrast, the CoCS model (Fig. 5f) yielded significantly larger apparent elastic moduli for soft and intermediate substrate stiffnesses, but similar moduli on stiff substrates. Moreover, when analyzed with the CoCS model, cell moduli did not depend on substrate stiffness, remaining around 100 Pa on all substrates (Fig. 5f). These results suggested that the overall stiffness of microglial cells is independent of substrate stiffness, similarly to that of polyacrylamide beads.

To test if the observed behavior is specific to microglial cells or a more general phenomenon, we repeated these experiments with fibroblasts, which have previously been suggested to adapt their stiffness to that of their environment^9^. As in our microglia experiments, fibroblasts only showed the apparent softening on softer substrates when using standard Hertz fits but did not show any significant changes in stiffness when analyzed using the CoCS model (Supplementary Fig. 7a). Also, similar to the bead and microglia experiments, the CoCS model fitted the fibroblast data on soft and intermediate substrates significantly better than the Hertz model, while both models worked similarly well on stiff substrates (Supplementary Fig. 7b). Together, these data suggested that cells do not adapt their overall mechanical properties to substrate stiffness.

These results were confirmed for samples exhibiting a coat such as pericellular brushes found in some cell types^8,24,25^ using FEM simulations and PAA beads functionalized with a polyethylene glycol (PEG) layer (Supplementary Fig. 8). Furthermore, we confirmed that cell curvature, which changes with substrate stiffness and cellular traction forces, has no impact on the validity of the CoCS model (Supplementary Fig. 9). Taken together, while the standard Hertz model underestimates elastic moduli of samples on substrates which are as soft as or softer than the sample, the CoCS model returns correct elastic moduli independent of substrate stiffness.

### Blebbistatin reduces ‘soft substrate effect’

In previous reports, perturbations of actomyosin contractility were shown to prevent the apparent stiffening of cells on stiffer substrates^26–28^. When we treated cells with the myosin II inhibitor blebbistatin, elastic moduli of microglial cells significantly decreased by about 20% (Supplementary Fig. 10). In contrast to our control experiments, the measured apparent elastic moduli of treated cells were independent of substrate stiffness and similar for both models (Fig. 5h). Furthermore, substrate deformability was generally smaller than without treatment (Fig. 5e) and similar across all different substrates (Fig. 5g). As blebbistatin reduced the overall cortical stiffness of the cells, it increased the ratio of *E*_substrate_/*E*_cell_. Hence, our data suggest that blebbistatin treatment increased the accuracy of the Hertz model on soft substrates because the contribution of *δ*_substrate_ to the measured total indentation decreased.

## Discussion

Here we show that, in AFM indentation measurements, the force exerted on a cell is transmitted to the soft substrate underneath, causing its deformation (Figs. 1, 2). The commonly used Hertz model therefore underestimates cortical cell stiffness on soft substrates with *E*_substrate_/*E*_cell_ ≲ 1 but converges towards correct values on stiffer substrates with *E*_substrate_/*E*_cell_ ≫ 1 (Figs. 3–6). To account for this ‘soft substrate effect’, we here developed the ‘composite cell-substrate model’ (‘CoCS’ model), which returns correct apparent elastic moduli independently of substrate stiffness, indentation depth *δ* (Supplementary Figs. 6, 7), pericellular coat (Supplementary Fig. 8), and cell curvature (Supplementary Fig. 9). The CoCS model does not require any knowledge about the cell-substrate geometry, and it can be implemented in any standard AFM indentation measurement.

**Figure 6:**
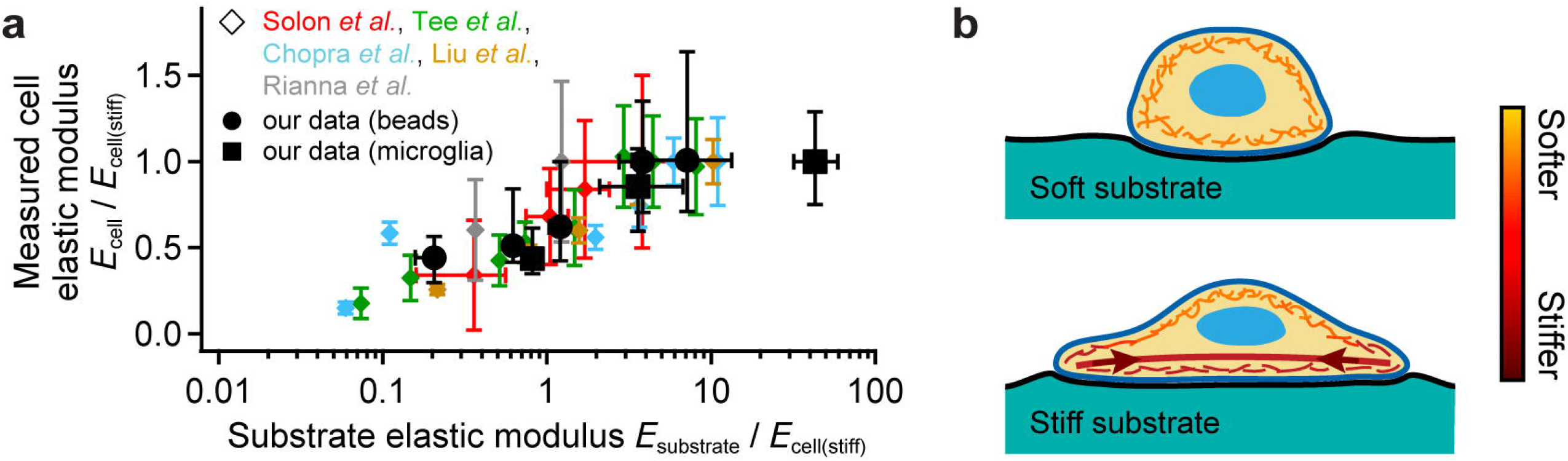
Comparison of normalized published and current data analyzed by the Hertz model and hypothesis. **(a)** Cell elastic moduli *vs*. substrate elastic moduli normalized by the respective cell elastic modulus on the stiffest gel used, *E*_cell(stiff)_. Note that data points collapse on a similar functional form. Data for various cell types from references^9–13^ (diamond symbols) and data for beads and cells from this study (circles and squares, respectively). Data points and error bars represent average and standard deviation or median and quartiles, respectively. **(b)** Schematic of force propagation in cells cultured on deformable substrates. On soft substrates (top), traction forces are small. On stiffer substrates (bottom), traction forces (arrows) generated mostly be ventral stress fibers^37^ (thick lines) increase with increasing substrate stiffness. These stress fibers undergo stress-stiffening and thus become stiffer with larger forces. These forces may be at least partly transmitted to the actomyosin cortex (thin fibers) but are dissipated with increasing distance from the stress fibers (illustrated by color going from red to orange). Hence, away from the stress fibers, the actin cortex does not stiffen significantly despite an increase in traction forces, as shown here by AFM measurements.

Previous reports using a Hertz model-based analysis of AFM indentation data suggested that the stiffness of cells increases with substrate stiffness^9–13^. We made similar observations when analyzing our own AFM data using the standard Hertz model (Fig. 6a). However, when correcting for the ‘soft substrate effect’ using the CoCS model, elastic moduli of both polyacrylamide beads and the cells remained largely constant and independent of substrate stiffness.

Microglial cells and fibroblasts spread more and exert higher traction forces as the substrate’s stiffness increases^29–31^, as confirmed in this study (Supplementary Fig. 9a-h). The current conceptual model explaining cortical cell stiffness-sensitivity to substrate mechanics hypothesizes that, as actomyosin-based traction forces of cells increase on stiffer substrates, the entire actin cytoskeleton stress-stiffens^9–13,26–28,32–36^. However, traction forces in two-dimensional cultures are mainly generated by ventral stress fibers^37^ rather than by the cortical actin network. While it is likely that stress fibers are coupled to the actin cortex, the lack of cortical stiffening in cells cultured on stiffer substrates (Fig. 5, Supplementary Fig. 7a) suggests that cellular traction forces are dissipated with increasing distance from the stress fibres, and that the distant actin cortex itself does not stress-stiffen on stiffer substrates (Fig. 6b).

In previous studies and our current work, blocking myosin II by blebbistatin abolished the apparent stiffening of cells on stiffer substrates when using the standard Hertz model. These findings were explained by the loss of contractility-driven stress stiffening of the actin cytoskeleton^10, 26–28^. However, blebbistatin does not only decrease the contractility of actin fibers (*i.e*., stress stiffening) but it also reduces the cell’s ‘base’ elastic modulus. Myosin II functions both as a motor protein and as a cross-linker^38^. As blebbistatin blocks myosin II in a detached state^39^, it leads to a decrease in cross-linking of the actin cortex. Because the elastic modulus of a polymer network such as the actin cortex non-linearly scales with the amount of cross-linking^40^, blebbistatin application leads to a global softening of the actin cortex, irrespective of traction forces, and thus to an increase in the *E*_substrate_/*E*_cell_ ratio. Hence, the ‘soft substrate effect’ is significantly reduced (Fig. 3e) and the Hertz model more accurate, providing an alternative explanation for why blebbistatin-treated cells do not seem to ‘soften’ on softer substrates (Fig. 5f).

When cells are cultured on stiff substrates, substrate effects in AFM measurements can be avoided by limiting the indentation depth to less than ∼10% of the sample height^41,42^. Importantly, this is not the case when cells are grown on soft substrates. Depending on the ratio of *E*_substrate_/*E*_sample_, samples may be pushed into the substrate even at very low forces, suggesting that this effect cannot be experimentally avoided when using AFM or any other nanoindentation approaches. However, the CoCS model provides a straight-forward tool to correct for this ‘soft substrate effect’, thus enabling AFM-based cell mechanics measurements on substrates with physiologically relevant stiffnesses. It is also likely applicable to cells within soft biological tissues, thus widening the scope and accuracy of AFM-based cell stiffness measurements.

## Acknowledgements

The authors would like to thank Paul Janmey, Ben Fabry, and Ulrich Schwarz for critical discussions and comments on the manuscript, and Alex Winkel (JPK) for technical support. NIH3T3 cells were a kind gift from William Colledge. We acknowledge funding from the German Science Foundation (DFG grant RH 147/1-1 to J.R., EXC 1003 CiM to T.B.), the Herchel Smith Foundation (postdoctoral fellowship to A.D.), the Royal Society (University Research Fellowship to K.J.C.), the UK EPSRC (Programme grant EP/P030017/1 to M.C.G.), the Human Frontier Science Program (HFSP grant RGP0018/2017 to T.B.), the European Research Council (Consolidator Grants 772798 to K.J.C., 771201 to T.B., 647186 to G.C., and 772426 to K.F.), and the UK BBSRC (Equipment grant BB/R000042/1 to G.C., and Research Project grant BB/N006402/1 to K.F.).

## Author contributions

J.R. and K.F. conceived the study; J.R. conducted all AFM experiments, analyzed all AFM data and developed the model, A.D. conducted all optical imaging and traction force microscopy experiments and analyzed data, B.W. and T.B. custom-designed polyacrylamide beads, N.M.K. and M.C.G. conducted ERISM measurements, K.J.C. helped with imaging and data analysis, G.C. helped with AFM experiments, all authors discussed the study, J.R. and K.F. wrote the paper with contributions from all co-authors.

## Competing financial interests

The authors declare no competing financial interests.

## Data and code availability

The data underlying this study are available from the authors upon reasonable request. Codes used for processing of AFM and confocal laser scanning microscopy raw data can be found at https://github.com/FranzeLab/AFM-data-analysis-and-processing/tree/master/Cell%20stiffness. Comsol models can be found at https://doi.org/10.6084/m9.figshare.10731869 and AFM force-distance curves raw data at https://doi.org/10.6084/m9.figshare.10732415.

## Additional information

Supplementary information is available in the online version of the paper.

## Methods

### Substrate preparation

Deformable PAA gel substrates as described previously^1,31,43^, Briefly, cover slips were glued into custom-made petri dishes, cleaned and silanized with (3-aminopropyl)trimethoxysilane (APTMS; unless otherwise stated, all chemicals from Sigma-Aldrich, Dorset, UK) for 3 min (minutes), treated with glutaraldehyde (diluted 1:10) for 30 min. Gel premixes were made by thoroughly mixing 440 μL of 40% acrylamide, 60 μL of 100% hydroxyl-acrylamide, and 250 μL of 2% Bis-acrylamide. The premix was then mixed with PBS at ratios between 40 μL to 460 μL and 150 μL to 350 μL to achieve gel stiffness between 50 Pa and 20 kPa. Polymerization was initialized by adding 0.3% (v/v) N,N,N’,N’-tetramethylethylenediamine (TEMED) and 0.1% (w/v) ammonium persulfate (APS) and 20 μL of the solution (giving about 100 μm gel thickness) were covered with cover slips, which were made hydrophobic with RainX (Shell Car Care International Ltd, UK) for 10 min beforehand. After at least 20 min top cover slips were removed, washed 2x with PBS, sterilized under UV light for 30 min, functionalized with either 100 μg/mL poly-D-lysine overnight for microglia cells or with 0.2 mg/mL fibronectin (FC010, Merck, 1:5 in PBS) for 2h at 37°C for fibroblasts, and washed 2x with PBS.

### PAA bead preparation

An AH-mix was produced by mixing 100 μL of 40% acrylamide with 13 μL of 97 % N-Hydroxyethyl acrylamide. Then, 50 μL of 2% Bis-acrylamide were added to 100 μL of AH-mix (ABH-mix). For the pre-bead-solution, first 100 μL of ABH-mix and 325 μL of PBS were mixed. This mixture was degassed for 10 min before adding 75 μL of 10% APS. The pH value of the pre-bead-solution was neutralized by adding 2.25 μL of 6M NaOH. An emulsion was generated by injecting 50 μL of pre-bead-solution in 500 μL of n-hexane with 3% Span^®^80 (Sigma-Aldrich) using a 100 μL Hamilton Syringe. After discarding the supernatant, the polymerization of the emulsion was initialized by adding 1.5 μL TEMED and keeping the emulsion at 85 °C for 10 min. After polymerization was finished, the beads were washed with n-hexane and transferred into 500 μL PBS. The elastic PAA beads were fluorescently labeled by preparing an ATTO488-solution of 1 mg ATTO 488 NHS-Ester (ATTO-TEC) in 200 μL Dimethylsulfoxide and adding 6 μL ATTO488-solution to the PAA beads in PBS. After 3 hours, the PAA beads were washed by centrifugation at a relative centrifugal force of 600 for 5 min. The supernatant was discarded and replaced by fresh PBS. The labeling and washing procedure was repeated three times. PAA beads were immobilized on the gel substrates by coating the substrates for 2 h with Cell-Tak (Corning Cat No 354240, 1:25 in PBS) and incubating the bead solution overnight at 4°C. By monitoring the beads in fluorescence microscopy during the AFM measurement it was ensured that beads were rigidly bound to the substrate. The strong adhesion resulted in a finite contact area between bead and substrate rather than a point contact (Fig. 4a and c, insets), making the bead-substrate contact analogous to a cell adhered a substrate, although the beads did not have a half-spherical shape. To investigate the influence of a pericellular coat on cell stiffness measurements, PAA beads were functionalized with a PEG layer. Beads where prepared similar as described above, however, instead of the ABH mix, a mixture of 100 μL of 40% acrylamide 50 μL of 2% Bis-acrylamide and 0.8 μL of Acylic-Acid (Sigma Aldrich, Germany) was generated (ABA-mix). For the PEG coating a 20kDa PEG polymer with NH2 and COOH groups on either end was used (NH2-PEG20K-COOH, Sigma-Aldrich, Germany). The NH2 side of the PEG polymer was bound to the beads by first activating the carboxyl groups exposed on the surface of the beads via ECD/NHS, allowing efficient covalent binding of the PEG. The still exposed COOH group of the PEG polymer allows to create multiple layers of the PEG coating. Here we performed this step 3 times to get an effective length of 60kDa. Briefly, an activation mixture was prepared using 1 ml of a NaCL/MES mix (solving 195mg MES, 4-Morpholineethanesulfonic acid, and 292 mg NaCl in 10ml pure water), which was added to 0.05 g EDC (N-(3-Dimethylaminopropyl)-N’-ethylcarbodiimide, Sigma-Aldrich, Germany). The coating was done in the following three step procedure: I) 1mL of the resulting solution was added to 11.5 mg NHS (N-Hydroxysuccinimide, Sigma-Aldrich, Germany) and vortexed. 500mL of this activation mix was added to the beads solution (500μL) and incubated for 15 minutes at room temperature. II) Immediately afterwards the beads where washed with PBS, and after a final centrifugation, 125mL of PBS was added after discarding the supernatant. 30μL of a NH2-PEG20K-COOH stock solution (500mg of NH2-PEG20K-COOH in 2500μL pure water) was added to the washed beads and incubated at room temperature. III) Finally, the beads where washed with PBS and filled up to 500μL total volume. The steps I)-III) where repeated three times.

### Culture preparation

All animal experiments of this study were conducted in accordance with the UK Animals (Scientific Procedures) Act (1986). Primary microglial cells were prepared from neonatal rat cerebral cortices as previously described^44^. Briefly, mixed glia cultures were prepared from neonatal rat cerebral cortices and cultured until they became confluent. Microglia and oligodendrocyte progenitor cells (OPCs) were then shaken-off at 320 rpm for 60 min and allowed to adhere for 20-25 min to uncoated culture dishes (Corning 430591), after which microglia but not OPCs adhere, which were then washed off. Fibroblasts were cultured in DMEM (with 10% FBS, 1% penicillin-streptomycin, glutamax). Microglial cells or fibroblast were then seeded on PAA substrates at a density of typically 10,000 cells/cm^2^ and cultured overnight. AFM measurements were performed in CO_2_-independent medium (Leibovitz L-15, w/o phenolred, with glutamax) at 37°C (using PetriDishHeater, JPK Instruments AG, Berlin, Germany). For inhibition of myosin, cells were incubated for 30 min in Leibovitz L-15 containing 20 μM blebbistatin (from stock solution 25 mM in DMSO) prior to measurements. For washout of blebbistatin, cells were washed three times and incubated for 30 min in fresh Leibovitz L-15 prior to measurement. For control measurements, 0.8 μL DMSO was added to 1mL of medium.

### Atomic force microscopy (AFM)

AFM measurements were performed on JPK Cellhesion 200 AFMs (JPK Instruments AG) installed either on a conventional (Axio Observer.A1, Carl Zeiss Ltd., Cambridge, UK) or a confocal optical microscope (see below). Tip-less AFM cantilevers (Arrow TL1, nominal spring constant *k* = 0.03 N/m, NanoWorld, Neuchâtel, Switzerland,) were calibrated using the thermal noise method^45^. Subsequently, monodisperse polystyrene micro-spheres (micro-particles GmbH, Berlin, Germany) with diameter 2*r* = 5 μm (PS/Q-R-KM153, Fig. 5) or 10 μm (Fig. 4) without or with fluorescence (diameter 5 μm, PS-FluoRed-Fi300, Fig. 1) where then glued to cantilevers (M-Bond 610, Micro-Measurements, Raleigh, NC, USA, agent and adhesive mixed at 1:1.3 weight ratio, cured at 80°C overnight). The sizes of the cantilever probes were chosen small to reduce influences by the limits of the Hertz model.

All AFM data were recorded at 1 kHz and subsequently filtered to 100 Hz sampling rates using binomial smoothing. For recording force *vs*. indentation data, the cantilever was positioned visually over at least three different, arbitrarily chosen positions above the cell or bead center identified by phase contrast microscopy^46,47^ and then approached at 5 μm/s until reaching a force set point of 500 pN for microglia cells, 1.5 nN for fibroblast, and 25 nN for beads. The force set point values were chosen to maximize signal to noise ratios while avoiding an influence of the finite cell height^41,42^. While taking more measurements across a cell would likely reduce the spread of the data, it would not change the results of our analysis, as the same force-distance curves were analyzed with both Hertz and CoCS models.

As common in AFM data analysis^48^, force *F* and tip indentation *δ* were calculated form the cantilever deflection *d* using the cantilever spring constant *k* and the vertical cantilever position *z* using *F* = *k* · (*d* – *d*_0_) and *δ* = *z* – *z*_0_ – *d*, where *z*_0_ and *d*_0_ denote the vertical cantilever position and deflection at the point of contact of the tip with the cell, respectively. The point of contact was detected as the first point where the force increased by threefold the standard deviation above baseline^48^, and the force curve was fit with the respective fit model (Equations (1) and (5)) between contact point and force set point using LMA least-squares fitting. For measuring the substrate stiffness, force curves were recorded on the substrate next to the cells or beads with force set points according to the substrate stiffness to maintain a consistent maximum indentation of about 2 μm.

Most fitting procedures for AFM data do not take the curvature of cells into account and assume that their surface is flat. This simplification usually only introduces a small error. As shown in our simulations, the influence of the cell curvature on the measured elastic moduli is indeed small and has no effect on the applicability of the CoCS model (Supplementary Fig. 5). Hence, in our main analysis we did not account for the curvature of the cells. However, as the curvature of cells changes on substrates of different stiffness (Supplementary Fig. 9m-r), we corrected both the Hertz and CoCS models for the cell curvature in Supplementary Figure 9 by replacing 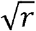 with 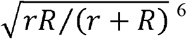. Even when accounting for the cell curvature (for details, see Supplementary Fig. 9m-p), the elastic moduli of the cells remained constant on all substrates when analyzed using the CoCS model, but appeared to ‘soften’ on softer substrates when analyzed using the Hertz model (Supplementary Fig. 9q and r).

### Confocal laser scanning microscopy

Combined AFM measurements and confocal microscopy were performed using a JPK Nanowizard AFM interfaced to a confocal laser scanning microscope (Olympus Fluoview FV1000, Olympus, Hamburg, Germany) equipped with a 40× silicon oil objective (NA 0.9, UPLSAPO, Olympus). For measuring the substrate displacement (Fig. 1a and b, Supplementary Fig. 1a and d, Supplementary Fig. 2a and b), *x-z* profiles were recorded through the cell center, while the cantilever was applied a constant force between 0.5 and 1.5 nN using the AFM’s force feedback. The substrate displacement was calculated by comparison of the two profiles using a modified cross correlation procedure to achieve sub-resolution accuracy (see Supplementary Fig. 1 for details). For confocal fluorescence microcopy, the PAA gels were fluorescently labeled by replacing 5 μL of the PBS with 1% (w/v) fluorescein O,O’-dimethacrylate in DMSO. Cells were incubated with 20 μM CellTracker Deep Red (Thermo Fisher Scientific) in serum-free medium for 30 min.

### Elastic resonator interference stress microscopy

ERISM substrates with an apparent stiffness of 3 kPa were fabricated as described previously^17^. A silicon chamber (surface area: 1.6 x 1.6 cm^2^, Ibidi) was applied to the ERISM substrate and the substrate surface was functionalized by incubating 1.5 mL of type 1 collagen (Collagen A, Biochrome) at pH 3.0-3.5 for one hour at 37 °C. After functionalization, the substrate was washed with cell culture medium (DMEM w/glutamax, 10% FCS, 1% P/S; Gibco). 3T3 fibroblasts (Sigma-Aldrich) were seeded at a density of 2,000 cells/cm^2^ and cultured for 24 hours. AFM indentation measurements were performed with a Nanosurf FlexAFM on an inverted microscope (Nikon Ti) fitted with a heated stage. A spherical glass bead with a diameter of 12 μm was glued to a cantilever (qp-SCONT, Nanosensors) with a force constant of 0.011 N/m (measured by the thermal-tuning method before attaching the bead). The cantilever deflection was calibrated by pushing the beaded cantilever against a rigid glass substrate using a known z-travel distance. Combined ERISM-AFM measurements were carried out in cell medium at 37 °C. First, maps of the vertical substrate deformation caused by the contractility of the cell were recorded by imaging the reflectance of the ERISM substrate at 201 different wavelengths between 550 and 750 nm as described previously^17^. Next, the AFM cantilever was lowered onto the center of the cell until a compression force of 0.5 nN was reached. The force was kept constant via a feedback loop while repeating the ERISM readout using a reduced wavelength range of 51 nm to accelerate the measurement (<5 s)^18^. The compression force was successively increased to 1.0 nN and 1.5 nN, respectively, and ERISM readout was repeated for both forces. A final ERISM measurement was performed after the AFM cantilever was fully retracted again to ensure cell contractility had not changed significantly over the course of AFM indentation. The substrate displacement under the cell caused by AFM indentation was obtained by subtracting the displacement map of the cell without AFM indentation from the displacement maps taken at the different AFM indentation forces. Filtered ERISM displacement maps (Gaussian blur with 1.6 μm bandwidth) were converted into stress maps using FEM as described in^17^.

### Modelling

Finite element calculations were performed using Comsol Multiphysics 4.1 (COMSOL AB, Stockholm, Sweden). Briefly, in an axisymmetric model cell and substrate were modelled as linear-elastic with Young’s modulus *E*_cell_ and *E*_substrate_, respectively, and incompressible (Poisson ratio *v* = 0.49), as generally assumed for living cells^15,24^ and hydrogels^21^. The contact between tip and cell was modeled as friction-less and the contact between cell and substrate was modelled as a direct mechanical contact. The mesh consisted of typically 10^4^ elements. The elastic displacement of cell and substrate was then calculated in response to a tip loading force between zero and 0.5 *E*_cell_ *r*^2^, resulting in a maximum indentation of typically *δ*_cell_ ≅ 0.4*r*.

### Traction force microcopy

PAA gels were prepared on imaging dishes (μ-Dish, Ibidi, Germany) as described above. Fluorescent nanoparticles (FluoSpheres carboxylate, 0.2□μm, crimson, Life Technologies, UK) were added to the PAA pre-mixes at a concentration of 0.2 % volume and were then placed in an ultrasonic bath for 30□s to separate the beads. After starting polymerization, the imaging dish was inverted to ensure that beads settled close to the gel surface. Fibroblasts or microglia were seeded onto PAA gels with shear storage moduli G′ of 100 Pa (microglia only), 1□kPa (fibroblasts and microglia) and 10□kPa (fibroblasts only). After 24□h, cells where imaged using an inverted microscope (Leica DMi8) at 37□°C and 5% CO2, equipped with a digital sCMOS camera (ORCA-Flash4.0, Hamamatsu Photonics), an EL6000 illuminator (Leica, Germany), and a 63× oil objective (NA1.4, Leica, Germany). Images were acquired using the Leica LAS X software. Fluorescence images of beads, and widefield images of cells were taken every 2□min. After the image acquisition, the culture media were exchanged with Trypsin-EDTA (Gibco) to detach cells from the gel. Reference images of fluorescent beads were taken 15□min after trypsinization. Traction stress maps were calculated for each frame using a TFM Software Package in MATLAB^49^. Traction stresses were averaged over time for each cell. Post-processing of the data and statistical analyses were done with a custom Python script. A detailed quantification of microglial traction forces on substrates of different stiffness can be found in a previous study^29^.

### Data processing and statistical analysis

AFM data and optical images were processed and analyzed in Igor Pro 6 (Wavemetrics, Portland, OR) using custom-written software. For measuring PAA beads and cells on substrates of different stiffness, for each bead / cell at least three force curves were recorded and analyzed, and their median values used. Presented values represent median unless otherwise stated. Box plots show median (band), quartiles (box), and standard error of median (notches), calculated as 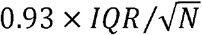 with *IQR* and *N* being interquartile-range and number of independent experiments, respectively.^50^ Goodness of fit was quantified using the adjusted coefficient of determination, 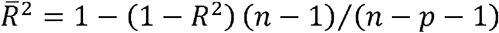, which accounts for the different number of fit parameters *p* with a number of data points *n* and coefficient of determination *R*^2^. As stiffness values followed log-normal distributions, statistical significance was tested using two-tailed Student’s *t*-tests (for two groups), two-tailed paired Student’s *t*-tests (for stiffness ratios), or oneway ANOVA followed by Tukey tests (for three or more groups) on logarithmized stiffness values. Deformability, 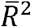, cell area, and mean traction stress values did not follow log-normal or normal distributions and were therefore tested for statistical significance using two-tailed Wilcoxon-Mann-Whitney *U* tests (for two groups) or one-way Kruskal-Wallis *H* test analysis of variance followed by Dunn-Holland-Wolfe tests (for three or more groups). Statistical significance was indicated using * for *P* < 0.05, ** for *P* < 0.01, and *** for *P* < 0.001, and “n.s.” for no statistical significant difference.

