## Supplement for "Cortical cell stiffness is independent of substrate mechanics"

### SUPPLEMENTARY FIGURES

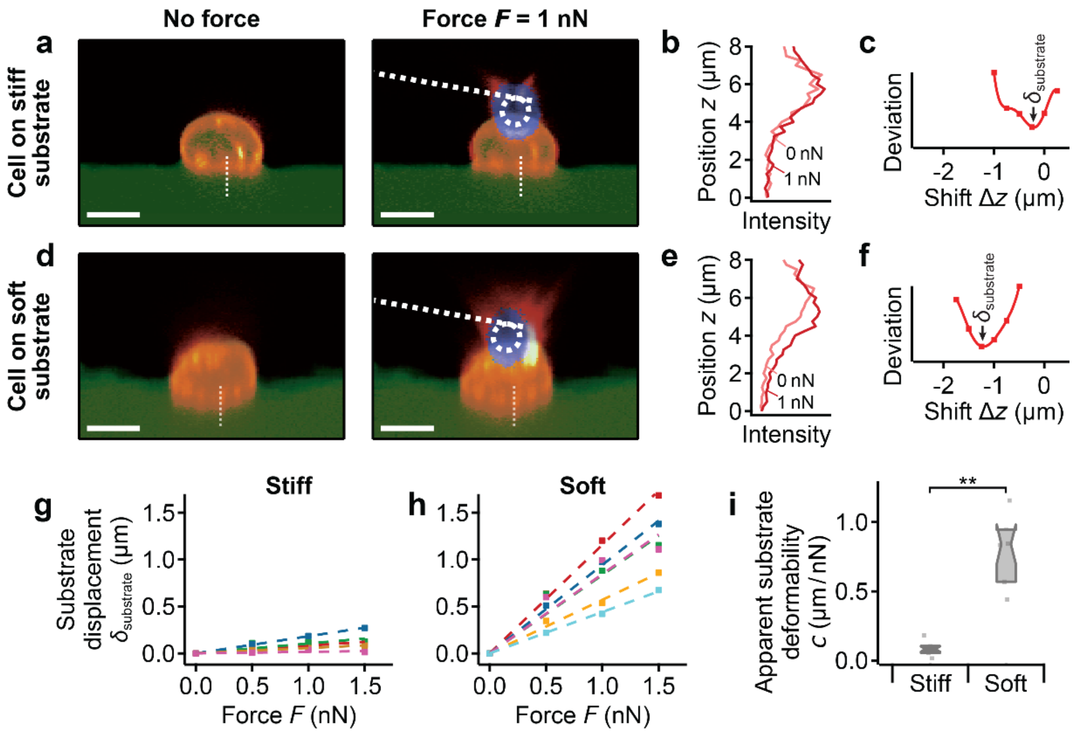


**Supplementary Fig. 1: Measuring substrate displacements with confocal microscopy. (a)** Confocal *z*-*x* profiles of a microglial cell on a stiff substrate without (left panel) and with the AFM applying a loading force (green: substrate; red: cell membrane; blue: AFM tip). **(b)** Profile of the red channel (location marked with dotted white lines in $x$-$z$-slices) without (light red traces) and with loading force $F = 1$ nN (dark red traces). **(c)** Deviation between the two profiles as a function of the vertical shift (markers) and fit of polynomial of degree 6 (continuous trace). The minimum indicates the measured substrate displacement $\delta_{\mathrm{substrate}}$. **(d)** Confocal *z*-*x* profiles, **(e)** profile of the red channel, and **(f)** deviation for a cell on a soft substrate. **(g, h)** Relationship between substrate displacements obtained from confocal images of multiple cells and the applied force *F* on **(g)** stiff and **(h)** soft substrates. **(i)** The deformability was significantly higher on soft compared to stiff substrates ($P = 0.0043$, Wilcoxon-Mann-Whitney *U* test). Animated versions of panels **(a)** and **(d)** are provided as movies online. Scale bars: 10 µm. Box plots show median (band), quartiles (box), standard error (notches), and data points (dots); $n = 5$ and 6 measurements from $N = 5$ cells each for the stiff and soft substrates, respectively. ** $P < 0.01$.

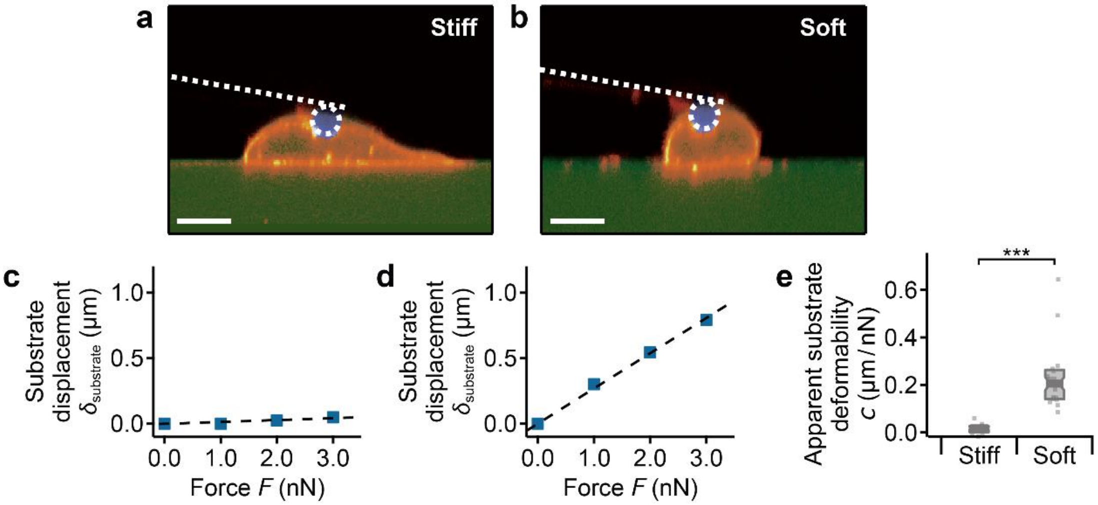


**Supplementary Fig. 2: Measuring substrate displacements with confocal microscopy for fibroblasts. (a, b)** Confocal *z-x* profiles of fibroblasts (orange) cultured on **(a)** stiff ($\approx 20$ kPa) and (**b)** soft ($\approx 300$ Pa) substrates (green). The AFM probe (blue) is applying a loading force of $F = 1$ nN on each cell. **(c, d)** Relationship between substrate displacements obtained from confocal images of the cells shown in **(a)** and **(b)** and the applied force *F* on **(c)** stiff and **(d)** soft substrates (see also Supplementary Fig. 1). **(e)** The deformability was significantly higher on soft compared to stiff substrates ($P = 3.3 \times 10^{-9}$, Wilcoxon-Mann-Whitney *U* test). Animated versions of panels **(a)** and **(d)** are provided as supplementary movies online. Scale bars: 10 µm. Box plots show median (band), quartiles (box), standard error (notches), and data points (dots); $n = 16$ measurements form $N = 8$ cells each for the stiff and soft substrates. *** $P < 0.001$.

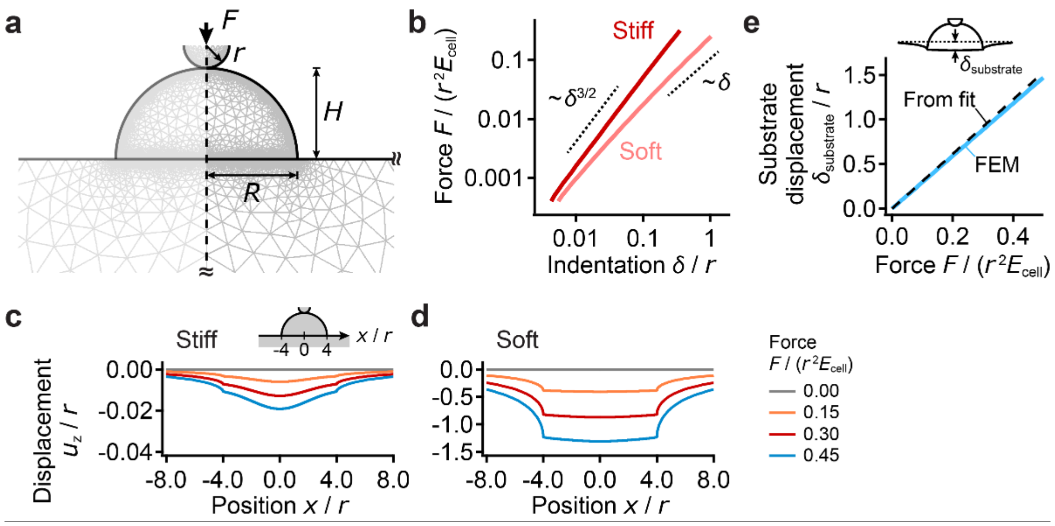


**Supplementary Fig. 3: Finite Element Model. (a)** Schematic of axisymmetric model for spherical tip with radius $r$ and loading force $F$ in contact with a cell of height $H$ and spreading radius $R$. The left half (grey) is mirrored for display but is not included in the model. **(b)** Force $F$ *vs.* indentation $\delta$ data from Figure 3c in log-log scale, generally showing $\delta^{3/2}$-dependency on stiff substrates and approaching $\delta^{3/2}$-dependency for small forces and linear $\delta$-dependency for higher forces on soft substrates. **(c)** Profiles of the resulting displacement (in units of the tip radius $r$) at the cell-substrate interface (inset shows the location where the profiles are taken, here the cell center is at $x = 0$ and the edges are at $x = \pm 4r$) for different loading forces (trace colors, in units of $r^2 E_{\text{cell}}$) for a stiff substrate and **(d)** for a soft substrate. **(e)** Calculated substrate displacement $\delta_{\text{substrate}}$ (measured at the cell center relative to the undeformed gel, see inset) *vs*. force (scaled in units of cell stiffness and tip radius) in comparison to the prediction from the CoCS model fit (dashed trace) for soft substrate. Parameters of calculation shown: cell height and radius $H = R = 4r$, $E_{\text{substrate}}/E_{\text{cell}} = 3$ ("stiff") and 0.03 ("soft").

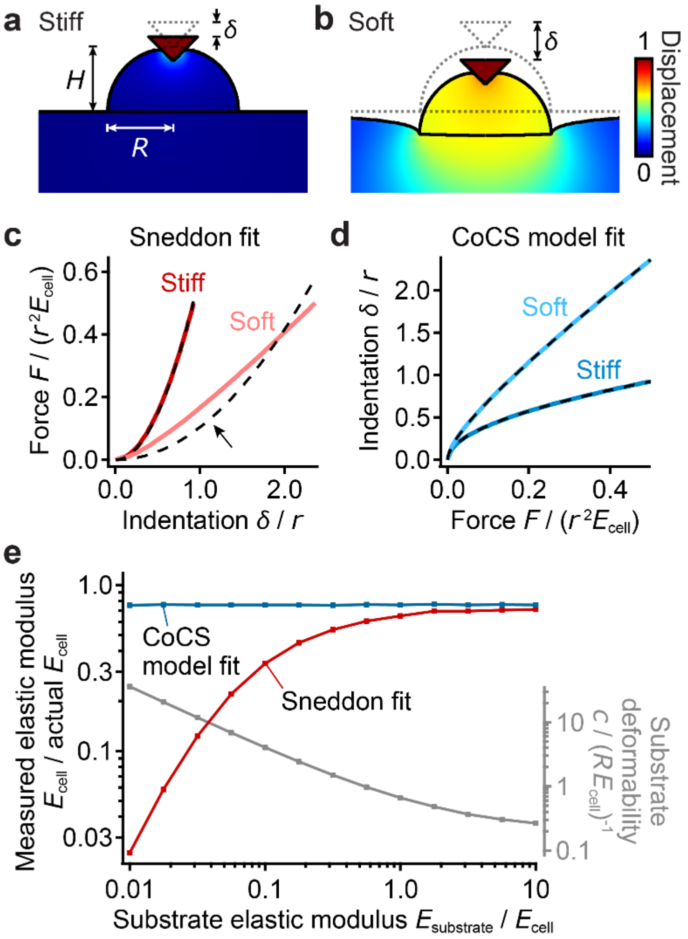


**Supplementary Fig. 4: Numerical validation for conical tips. (a, b)** Representative FEM results for cells on **(a)** stiff and **(b)** soft substrates for a force $F = 0.5\, r^2 E_{\mathrm{cell}}$. Color shows material displacement in units of tip displacement. $\delta$ indicates the measured total indentation relative to the undeformed state (dotted outlines). **(c)** Force $F$ *vs.* indentation $\delta$ for cells cultured on stiff and soft substrates analyzed with standard Sneddon model fits, $F(\delta) = 2/\pi \cdot E_{\mathrm{cell}}/(1-\nu_{\mathrm{cell}}^2) \cdot \tan\alpha \cdot \delta^2 \approx 8/3\pi \cdot E_{\mathrm{cell}} \cdot \tan\alpha \cdot \delta^2$ (dashed traces)[1]. The Sneddon model deviates from the data in measurements on soft substrates (arrow). **(d)** Indentation $\delta$ *vs.* force $F$ for cells on soft and stiff substrates with CoCS model fits, Equation (5) with $a = (8/3\pi \cdot E_{\mathrm{cell}} \cdot \tan\alpha)^{-1/2}$ and $b = 1/2$ (dashed traces). **(e)** Elastic moduli in units of the actual elastic moduli of the cells $E_{\mathrm{cell}}$ as a function of relative substrate stiffness as obtained by fitting force-indentation curves simulated by FEM using a standard Sneddon fit (red trace), or using the CoCS model fit (Equation (5), blue trace). Right axis shows substrate deformability obtained from the CoCS model fit (continuous gray trace). Parameters of calculations shown: cell height and radius $H = R = 4r$, $E_{\mathrm{substrate}}/E_{\mathrm{cell}} = 3$ ("stiff") and 0.03 ("soft").

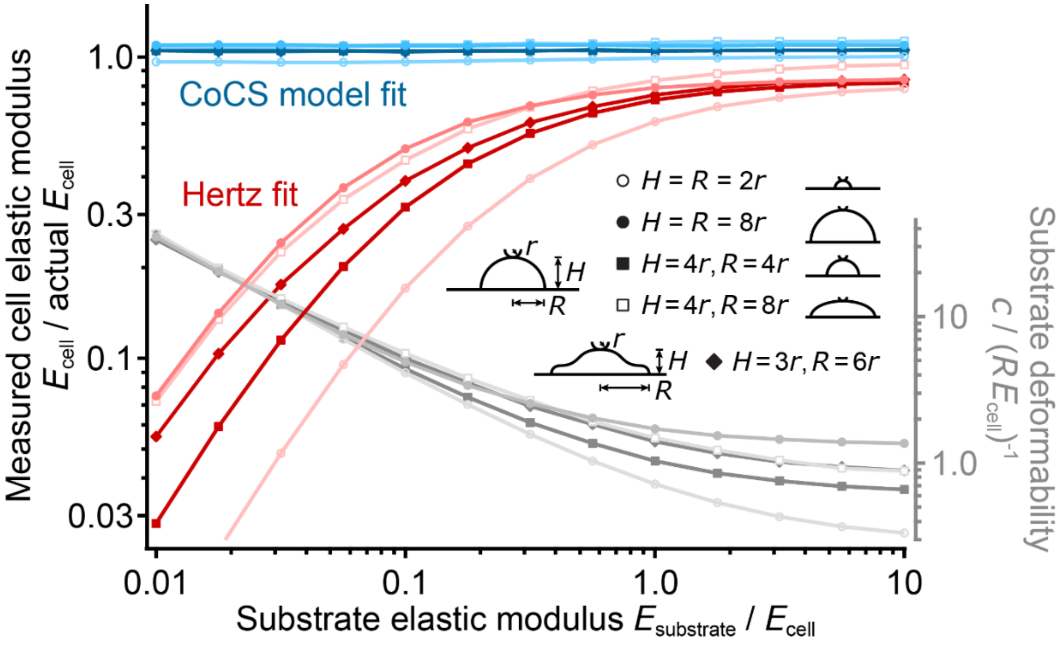


**Supplementary Fig. 5: Influence of the cell geometry on the measured stiffness.** Shown are elastic moduli $E_{\mathrm{cell}}$ obtained from force-indentation curves simulated by FEM and analyzed using standard Hertz model fits (Equation (1), red traces) and CoCS model fits (Equation (5), blue traces) as a function of the relative substrate stiffness for different cell geometries in terms of cell height $H$ (circles) and radius $R$ (squares) and for a non-spherical cell shape (diamonds). The right axis shows the substrate deformability obtained from CoCS model fits. The Hertz fits strongly depend on the cell geometry; hence correcting AFM data analyzed using the standard Hertz model for soft substrate effects would require precise knowledge of the cell's geometry and the substrate stiffness (see also [2,3]). In contrast, the CoCS model returns the correct cell stiffness independent of the cell geometry and substrate stiffness and without any prior assumptions about them. Thus, no prior knowledge about these quantities is required when fitting AFM data using the CoCS model.

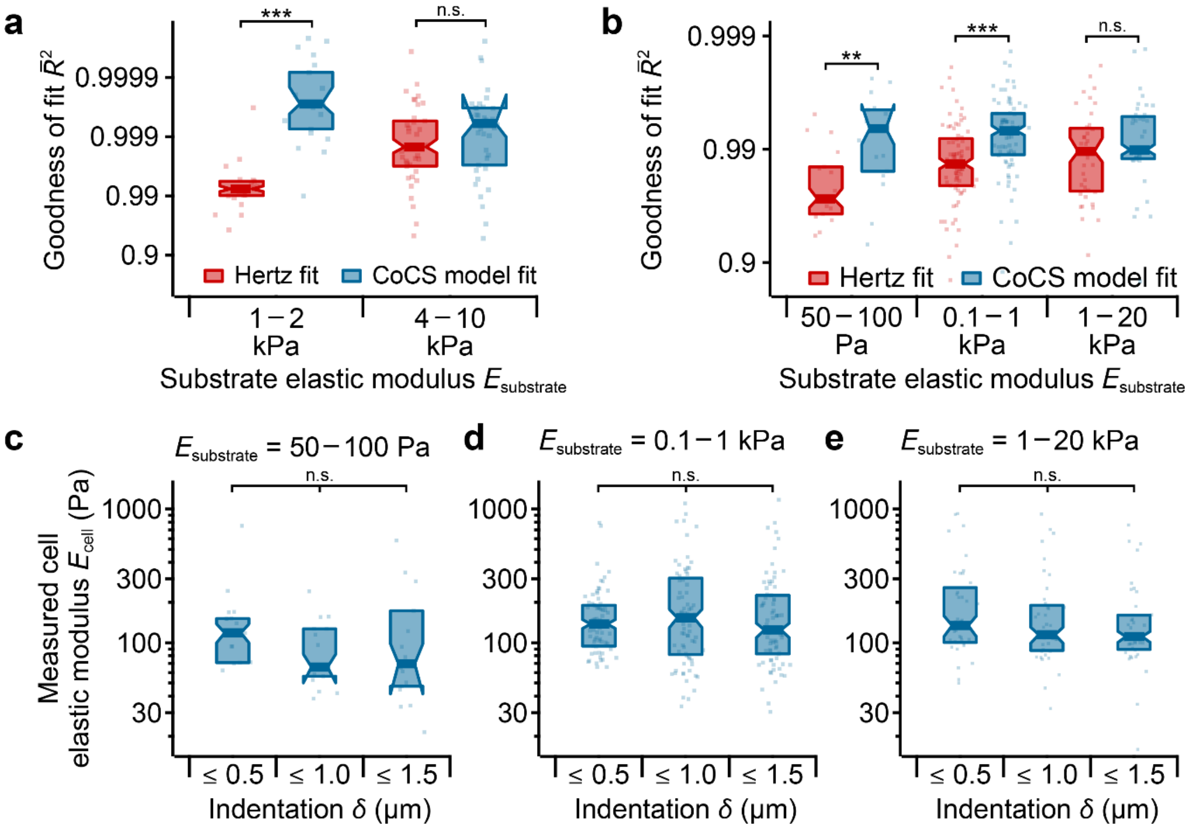


**Supplementary Fig. 6: Goodness of fit analysis.** Adjusted coefficient of determination $\bar{R}^2$ of Hertz and CoCS model fits for **(a)** the PAA bead data from Figure 4 and **(b)** the microglia data from Figure 5. The CoCS model fits AFM indentation data significantly better than the Hertz model for measurements of beads on soft substrates ($P = 6.8 \times 10^{-7}$, Wilcoxon-Mann-Whitney *U* test), but they fit similarly well on stiff substrates ($P = 0.21$). Likewise, for microglia cells, the CoCS model fit the experimental data significantly better than the Hertz model on soft and intermediate substrates ($P = 0.0069$ and $1.7 \times 10^{-5}$, respectively), while they fit similarly well on stiff substrates ($P = 0.20$). **(c-e)** Influence of the indentation depth $\delta$ on the measured elastic moduli of microglial cells (*cf.* Figure 5f) for **(c)** soft, **(d)** intermediate, and **(e)** stiff substrates. Apparent elastic moduli were similar (~$100$ Pa) for all indentation depths ($P = 0.88$, $0.52$, and $0.70$, respectively, one-way ANOVA) on all substrates, confirming self-consistency of the CoCS model. Box plots show median (band), quartiles (box), standard error (notches), and data points (dots); number of beads (a) $N = 21$ and $39$ for the soft and stiff substrates, respectively, and number of cells (b-e) $N = 17$, $74$, and $39$ for the soft, intermediate, and stiff substrates, respectively. ** $P < 0.01$, *** $P < 0.001$.

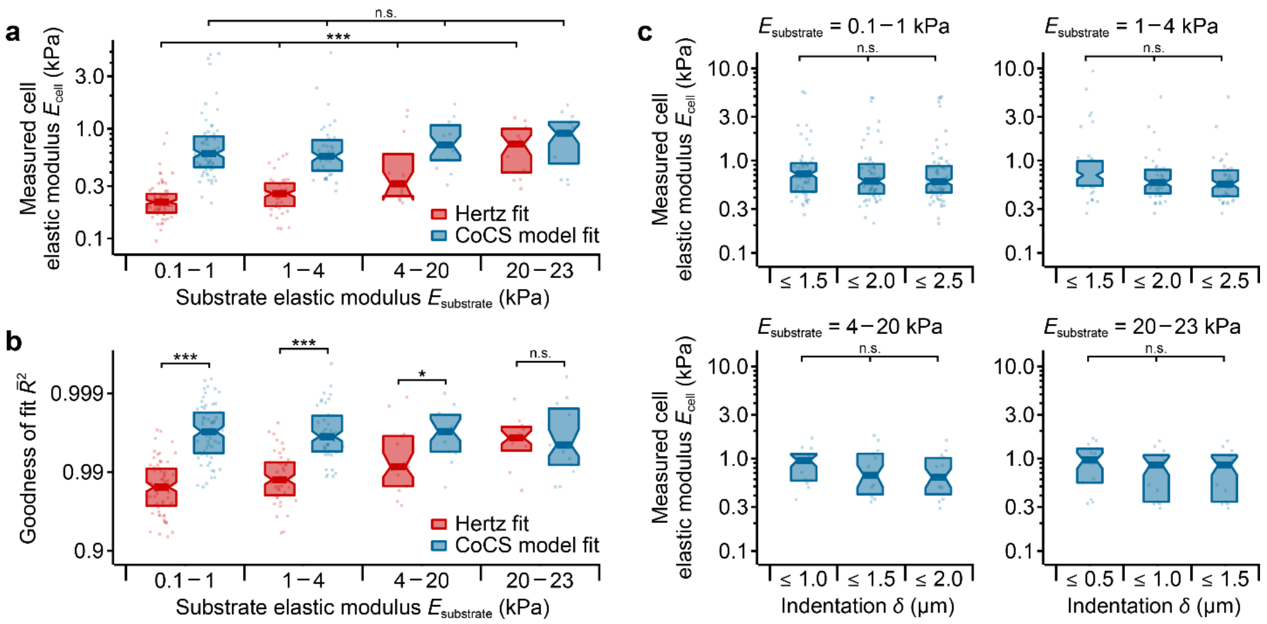


**Supplementary Fig. 7: Application of the CoCS model to AFM indentation measurements of fibroblasts. (a)** Apparent elastic moduli of live fibroblasts on substrates of different stiffnesses as obtained from standard Hertz fits (Equation (1), red), and from CoCS model fits (Equation (5), blue). As in microglial cell experiments (Figure 5), measured elastic moduli of fibroblasts remained constant on all substrates when analyzed using the CoCS model ($P = 0.7$, one way ANOVA), but appear to 'soften' on softer substrates when analyzed using the Hertz model ($P = 3.0 \times 10^{-12}$, one way ANOVA; $P \leq 0.004$, Tukey posthoc tests). **(b)** Adjusted coefficient of determination $\bar{R}^2$ of Hertz and CoCS model fits for the fibroblast data shown in (a). As for beads and microglial cells (Supplementary Fig. 6a, b), the CoCS model fitted AFM indentation data significantly better than the Hertz model for measurements of fibroblasts on soft and intermediate stiffness substrates ($P = 1.2 \times 10^{-15}$, $3.6 \times 10^{-9}$, and $0.029$, respectively, Wilcoxon-Mann-Whitney $U$ test), but both models fitted the data similarly well on stiff substrates ($P = 0.94$). **(c)** Influence of the indentation depth $\delta$ on the measured elastic moduli of fibroblasts. Apparent elastic moduli were similar (~$1$ kPa) for all indentation depths ($P = 0.81$, $0.49$, $0.72$, and $0.71$, respectively, one-way ANOVA) on all substrates, confirming self-consistency of the CoCS model also for fibroblasts. Box plots show median (band), quartiles (box), and standard error (notches); $N = 62$, $41$, $13$, and $13$ cells for the different substrate stiffness ranges. * $P < 0.05$, *** $P < 0.001$.

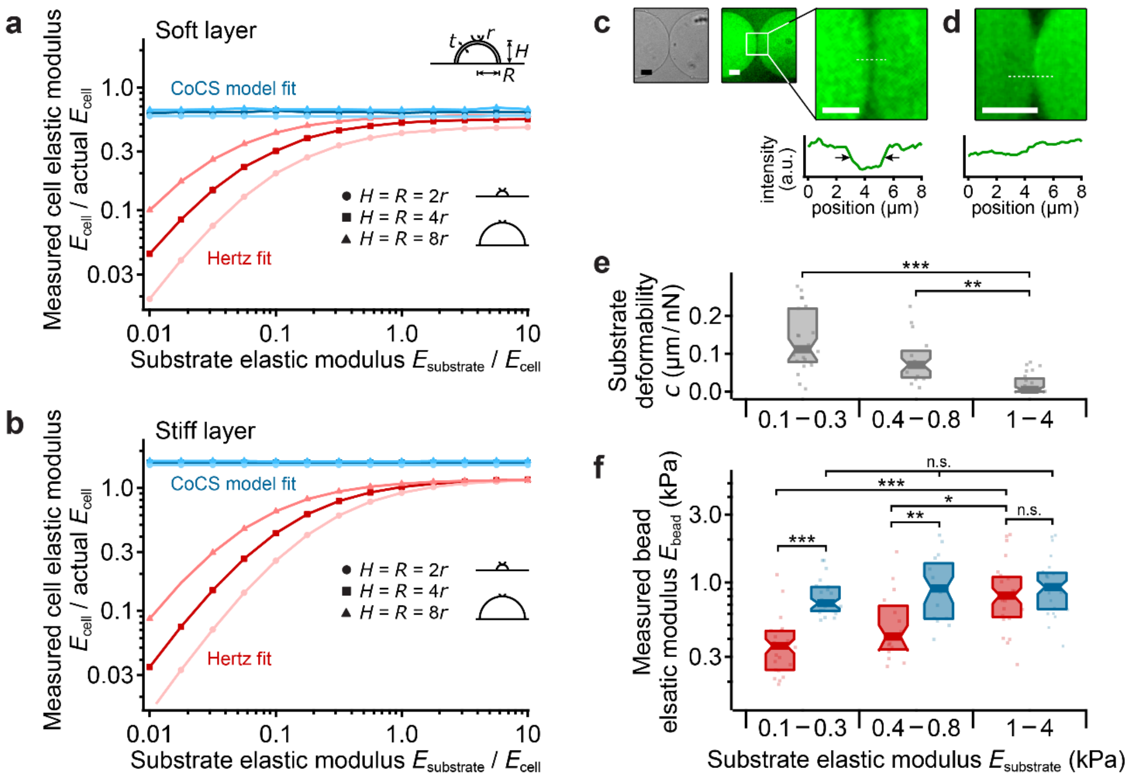


**Supplementary Fig. 8: Evaluating the influence of pericellular coats. (a, b)** Cell elastic moduli in units of the actual cell elastic modulus $E_{\mathrm{cell}}$ as a function of relative substrate elastic modulus as obtained from fitting force-indentation curves simulated by FEM using a standard Hertz fit (Equation (1), red trace), or using the CoCS model fit (Equation (5), blue trace) for **(a)** a soft elastic layer and **(b)** for a stiff elastic layer around the cell (see schematic in a) with thickness $t = 0.5r$ (corresponds to $\approx 1\ \mu\mathrm{m}$ for our experiments) and stiffness $0.5E_{\mathrm{cell}}$ (a) and $2E_{\mathrm{cell}}$ (b). As for the model without coat, CoCS and Hertz fit yielded the same stiffness for $E_{\mathrm{substrate}}/E_{\mathrm{cell}} \gg 1$. However, a soft or stiff pericellular coat resulted in a general under- or overestimation of the cell stiffness, respectively, as the AFM effectively measures a mixture of coat and cell stiffness. Nevertheless, while the Hertz fit underestimated the cell stiffness for $E_{\mathrm{substrate}}/E_{\mathrm{cell}} \lesssim 1$, the CoCS model fit yielded the cell stiffness independently of the substrate stiffness. **(c)** Bright field (left) and confocal images (right) of fluorescent polyacrylamide beads with molecular brush coats. Although the beads are in direct contact, the fluorescence signal is separated by $\sim 2 - 3\ \mu\mathrm{m}$ (arrows) because of the presence of non-fluorescent brush layers. **(d)** For comparison, beads without molecular brush coats touch each other directly. Scale bars: $10\ \mu\mathrm{m}$ **(e)** Substrate deformability obtained from CoCS model fits and **(f)** measured elastic modulus of beads with

PEG layer on substrates of different stiffness obtained from Hertz fits (Equation (1), red), and from CoCS model fits (Equation (5), blue). Note that, although the beads appear generally softer, as for the uncoated beads (Figure 4), the bead stiffness is independent of substrate stiffness when using the CoCS model fit but appears correlated with substrate stiffness when using standard Hertz fits. * $P < 0.05$, ** $P < 0.01$, *** $P < 0.001$.

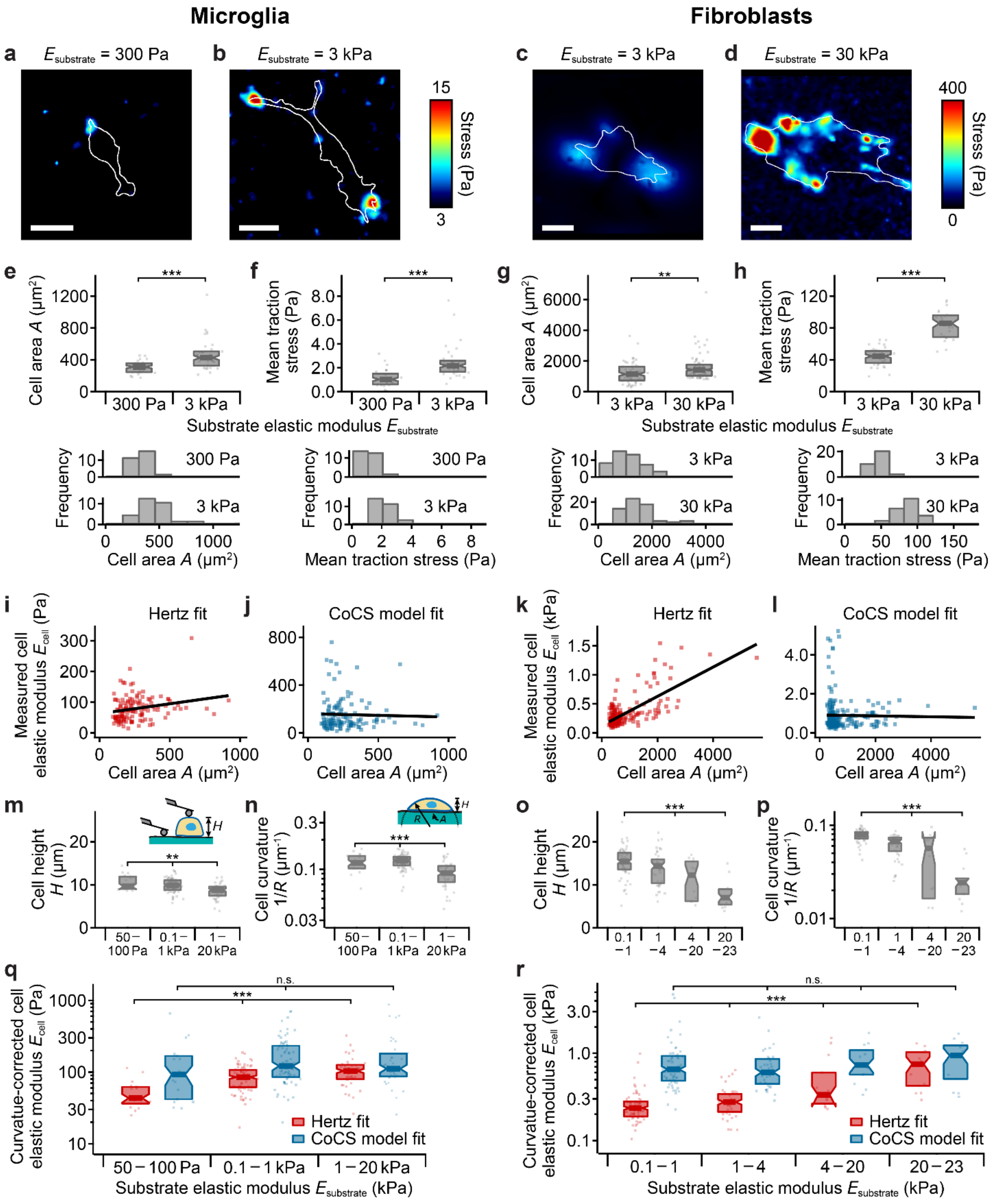


**Supplementary Fig. 9: Adaptation of microglial cells and fibroblasts to substrates of increasing stiffness. (a, b)** Traction force maps of microglial cells (white outlines) cultured on **(a)** soft and **(b)** stiff substrates and **(c, d)** of fibroblasts cultured on **(c)** soft and **(d)** stiff substrates. Cellular traction forces increase with increasing substrate stiffness. Both cell types showed the expected morphological phenotypes[4,5], with more spherical shapes on softer substrates (left) and well-spread morphologies with distinct protrusions on stiffer substrates (right). **(e)** Box plots (top) and histograms (bottom) of cell area and **(f)** mean traction stress of microglial cells and **(g)** cell area and **(h)** mean traction stress of fibroblasts on substrates of different stiffnesses, showing significant increases in both area and mean traction stress on stiffer substrates for both microglia and fibroblasts ($P = 2.1 \times 10^{-7}$, $4.1 \times 10^{-5}$, $0.0017$, and $8.4 \times 10^{-11}$, Wilcoxon-Mann-Whitney *U* tests). **(i-l)** Scatter plots of cell area *vs.* cell stiffness obtained from Hertz fits **(i, k)** and CoCS model fits **(j, l)**. For the Hertz fits, elastic moduli and area were highly correlated with cell area. Linear regressions (black lines) had slopes significantly different from zero ($P = 0.0070$ for microglia **(i)** and $P = 2.3 \times 10^{-32}$ for fibroblasts **(k)**, two-tailed *t*-test); Pearson's correlation coefficient $\rho = 0.23 \pm 0.09$ for microglia and $\rho = 0.70 \pm 0.05$ for fibroblasts. However, for the CoCS model fits, cell elastic moduli and area were not correlated ($\rho = 0.0 \pm 0.1$ and $P = 0.74$ for microglia; $\rho = 0.0 \pm 0.1$; $P = 0.79$ for fibroblasts). **(m-p)** Cell height $H$ and cell curvature $1/R$ for microglial cells (**m** and **n**, respectively) and fibroblasts **(o, p)** on substrates of different stiffnesses. Cell height was determined from AFM force-distance curves recorded on the cell and on the substrate next to the cell (see schematic in m). Cell curvature $1/R = 2H/(H^2 + A/\pi)$ was estimated assuming the cells having a spherical cap shape with cell height $H$ and cell area $A$ (see schematic in n). For both cell types, cell height and curvature significantly depended on substrate stiffness ($P = 0.0023$, $1.9 \times 10^{-9}$, $1.2 \times 10^{-9}$, $1.8 \times 10^{-9}$, one way ANOVA). **(q, r)** Cell curvature-corrected apparent elastic moduli for microglial cells **(q)** and fibroblasts **(r)** on substrates of different stiffnesses as obtained from standard Hertz fits (Equation (1), red), and from CoCS model fits (Equation (5), blue). To correct for cell curvature, in both models the elastic modulus was corrected by a factor of $\sqrt{1 + r/R}$ [6]. When accounting for cell curvature, the elastic moduli of microglia and fibroblasts remained constant on all substrates when analyzed using the CoCS model ($P = 0.27$, $0.76$, one way ANOVA), but appeared to 'soften' on softer substrates when analyzed using the Hertz model ($P = 9.5 \times 10^{-7}$, $1.6 \times 10^{-11}$, one way ANOVA). Scale bars: 20 µm (a-d). Box plots show median (band), quartiles (box), standard error (notches), and data points (dots); number of cells (e, f) $N = 30$ and $34$, (g) $57$, $53$, and $71$, and (h) $36$ and $24$, for the soft, intermediate and stiff substrates, (i, j) $131$ and (k, l) $227$. ** $P < 0.01$, *** $P < 0.001$.

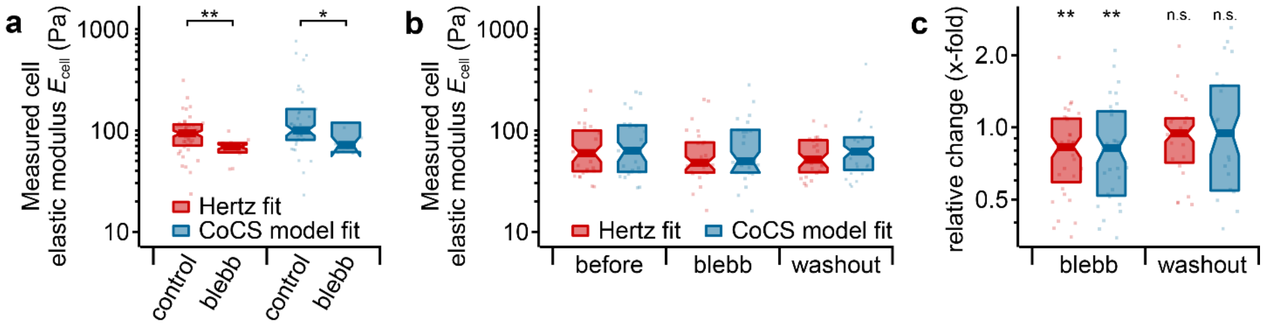


**Supplementary Fig. 10: Blebbistatin application to microglia cells on elastic substrates. (a)** Stiffness of microglia before and after treatment with blebbistatin on stiff substrates (same data as shown in Figure 5f and h), as obtained from standard Hertz fits (Equation (1), red) and from CoCS model fits (Equation (5), blue). Cortical stiffness decreased significantly after blebbistatin application ($P \leq 0.018$, paired *t*-tests). **(b)** Absolute values and **(c)** relative changes of cortical microglia stiffness before and after treatment with blebbistatin and after washout, summarized over different substrate stiffnesses blue), showing a relative decrease in cortical stiffness of ~20% after treatment with blebbistatin ($P \leq 0.006$, paired *t*-tests) and a recovery to base values after washout ($P \geq 0.24$, paired *t*-tests). Box plots show median (band), quartiles (box), and standard error (notches); number of cells (a) $N = 39$ and 12 for control and with blebbistatin, respectively, and (b,c) $N = 36$, 36, and 31 cells for before and after treatment and after washout, respectively. * $P < 0.05$, ** $P < 0.01$.

### SUPPLEMENTARY DISCUSSION

Given that it is established that cell function is critically regulated by substrate mechanics[7-11], soft substrates mimicking the mechanical properties of the physiological cell environment are widely used. Cell stiffness measurements on these substrates are crucial to understanding mechanical interactions of cells with their environment. However, measurements of cortical cell stiffness on deformable substrates are challenging, and current suggested correction methods to account for substrate mechanical properties are rather complex[2,3].

We here present a straightforward approach for estimating the apparent elastic moduli of cells on deformable substrates from AFM force-indentation curves, which can be applied without any prior knowledge of cell morphology (Supplementary Fig. 5) or the need for hardware modifications. Motivated by simple analytical considerations, we used ground-truth data from numerical simulations and experimental data from soft elastic polyacrylamide beads on polyacrylamide substrates to validate our method, which we termed composite cell-substrate (CoCS) model.

The substrate deformability $c$ obtained from CoCS model fits directly depends on the substrate's elastic modulus, as predicted by the analytical model, Equation (6), and by FEM (Fig. 3e). Hence, $c$ can also be used to estimate the substrate's elastic modulus $E_{\mathrm{substrate}} \approx 1/(Rc)$, obtained by rearranging Equation (6). Using a typical contact radius $R \approx 20\ \mu\mathrm{m}$ (see Fig. 4a, c), our experiments yielded elastic moduli of $E_{\mathrm{substrate}} \approx 1.0 \pm 0.1\ \mathrm{kPa}$ for the soft and $7 \pm 1\ \mathrm{kPa}$ for the stiff substrates, in reasonable agreement with the actual substrate elastic moduli measured directly of $1.4 \pm 0.1\ \mathrm{kPa}$ and $9 \pm 2\ \mathrm{kPa}$, respectively (Fig. 4e).

Cell stiffness characterizes the resistance of cells to deformation in response to forces. In the case of small externally applied forces as in the current study, the deformation is largely determined by peripheral cellular structures. Blebbistatin significantly reduced the apparent elastic moduli of the cells (Fig. 5h, Supplementary Fig. 8), suggesting a significant contribution of the actomyosin cortex to the measured values. Thick pericellular brushes found in some cell types will also contribute to apparent elastic moduli measured by AFM[12-14]. To investigate the predictions of the CoCS model for samples exhibiting a pericellular coat, we added a layer representing a pericellular coat to the FEM simulations. Also here, the CoCS model yielded the cell stiffness independently of substrate stiffness, while the Hertz fit underestimated the cell stiffness for soft substrates (Supplementary Fig. 9a, b). Furthermore, we functionalized PAA beads with a polyethylene glycol (PEG) layer mimicking a pericellular coat (Supplementary Fig. 9c). As for the uncoated beads (Fig. 4) and FEM simulations, elastic moduli were independent of substrate stiffness when using the CoCS model but correlated with substrate stiffness when using standard Hertz fits (Supplementary Fig. 9). Taken together, the CoCS model works well also for cells with pericellular coats, returning moduli independent of substrate stiffness (Supplementary Fig. 9), and it can be combined in the future with other models[15] to disentangle the contributions of the coat and the cells to the measurements.

Cortical cell stiffness is cell type-specific, may depend on chemical signaling[16], and change during pathological processes such as cancer metastasis[17]. Similar to chemical signals, mechanical signals may impact cell behavior *in vitro* as well as *in vivo*[18,19]. For example, an increase in substrate stiffness leads to an increase in cellular traction forces and in cell spreading[20,21] (Supplementary Fig. 10a-h). Previous reports using a Hertz model-based analysis of AFM indentation data also suggested that the stiffness of cells cultured on soft polyacrylamide gels first increases with increasing substrate stiffness and then plateaus. This behavior has been described for a variety of cell types including fibroblasts[5], human mesenchymal stem cells[22], aortic valve interstitial cells[23], thyroid cells[24], and cardiac myocytes[25]. Our results suggest, however, that the observed apparent softening of cells on soft substrates is due to the overestimation of their indentation when using the standard Hertz model to analyze AFM data. On substrates as soft as or softer than the cells, forces applied to cells lead to significant deformations of the underlying substrate, which can no longer be neglected. Accounting for this ‘soft substrate effect’ by using the CoCS model developed in this study revealed that the cortical cell stiffness is actually largely independent of substrate mechanics.
